# Marizomib suppresses triple-negative breast cancer via proteasome and oxidative phosphorylation inhibition

**DOI:** 10.1101/801720

**Authors:** Prahlad V. Raninga, Andy Lee, Debottam Sinha, Lan-feng Dong, Keshava K. Datta, Xue Lu, Priyakshi Kalita-de Croft, Mriga Dutt, Michelle M. Hill, Normand Pouliot, Harsha Gowda, Murugan Kalimutho, Jiri Neuzil, Kum Kum Khanna

**Author notes:** To whom correspondence should be addressed: Prof Kum Kum Khanna, Dr Prahlad Raninga.

## Abstract

Lacking effective targeted therapies, triple-negative breast cancer (TNBCs) is highly aggressive with development of metastasis especially brain, and remains clinically challenging breast cancer subtype to treat. Despite the survival dependency on the proteasome pathway genes, FDA-approved proteasome inhibitors induced minimal clinical response in breast cancer patients due to weak proteasome inhibition. Here, we show that a potent proteasome inhibitor Marizomib (Mzb) inhibits multiple proteasome catalytic activities and induces a better anti-tumor response in TNBC cell lines and patient-derived xenografts alone and in combination with the standard-of-care chemotherapy. Mechanistically, Mzb inhibits oxidative phosphorylation (OXPHOS) via PGC-1α suppression in conjunction with proteasome inhibition in TNBC cells. Mzb reduces lung and brain metastases by reducing the number of circulating tumor cells and the expression of multiple genes involved in the epithelial-to-mesenchymal transition. Furthermore, Mzb-induced OXPHOS inhibition upregulates glycolysis to meet the energetic demands of TNBC cells and, hence, combined inhibition of glycolysis with Mzb exposure leads to a synergistic anti-cancer activity. Collectively, our data provide a strong rationale for a clinical evaluation of Mzb in primary and metastatic TNBC patients.

**One Sentence Summary:** Marizomib inhibits primary tumor growth, and also reduces lung and brain metastases in pre-clinical models of triple-negative breast cancer.

## Introduction

Triple-negative breast cancer (TNBCs), defined by their lack of estrogen (ER), progesterone (PR), and Her2 receptors, is a very aggressive subtype of breast cancer (BC) that typically affects young women and has a wide variability in patient outcomes. Although a majority of TNBC patients respond to initial chemotherapies, these tumors are prone to recur, metastasize (disseminate to the lungs and brain), and become resistant (1). TNBCs are enriched with poorly differentiated tumor-initiating cells, which are often responsible not only for tumor initiation but also for metastasis and chemoresistance. Thus, there is an urgent need to explore clinically viable options for treatment of primary and metastatic TNBCs.

Proteasome inhibitors have been shown to exert a significant anti-cancer activity in multiple cancer types including multiple myeloma and leukemia. A genome-wide siRNA screen identified the survival dependency of TNBC compared to other breast cancer subtypes on proteasome genes (2), suggesting that the proteasome complex represents a plausible therapeutic target in TNBCs due to disrupted proteostasis contributed by their inherent genomic instability (3). Although a first-generation FDA-approved proteasome inhibitor, bortezomib (Btz), and second-generation inhibitors including carfilzomib (Cfz) and ixazomib (Ixz) induced a significant anti-tumor activity in pre-clinical models (2,4), these agents have not demonstrated efficacy in clinical trials in solid tumors including TNBCs, either alone or in combination with other therapies (4). This could be due to weaker proteasome inhibitory activity of these drugs in solid tumors. Furthermore, as a single agent, Cfz showed no tumor suppressive effect in an MDA-MB-231 xenograft model (5).

The 26S proteasome is composed of a cylindrical 20S core that is capped by the regulatory 19S component (6). The 20S core contains three proteasome active-site subunits within its β rings including the β1 site (caspase-like), β2 site (trypsin-like), and β5 site (chemotrypsin-like) (7). While Btz, Cfz, and Ixz selectively inhibit the chemotrypsin-like proteasome activity (CT-L) (8), they fail to inhibit the trypsin-like (T-L) and caspase-like (CP-L) proteasome activity, which are known to degrade proteins (9). A recent study has shown that Btz/Cfz-induced CT-L proteasome activity inhibition resulted in upregulation of T-L and CP-L proteasome activities in Nrf1-dependent manner (10), which is a dominant mechanism of resistance to these drugs. Notably, selective inhibition of T-L (β2) proteasome activity sensitized TNBC cells to Btz/Cfz-induced apoptosis (10). Hence, co-inhibition of multiple active sites of the proteasome complex could increase the potency and anti-tumor activity of proteasome inhibitors in solid tumors including TNBCs.

Marizomib (Mzb), also known as NPI-0052, is an orally active, small molecule proteasome inhibitor derived from marine bacteria (11). Mzb has been shown to inhibit all three active sites (CT-L, T-L, and CP-L) of the proteasome in multiple myeloma and in solid tumors (8,12), thus has increased potency to be a multi-site antagonist of the proteasome complex compared to other peptide-based proteasome inhibitors. Based on this information, we hypothesized that Mzb, by simultaneously inhibiting multiple proteasome catalytic sites, will show enhanced anti-tumor activity in TNBCs compared to the previous generation compounds.

In this study, we examined the anti-cancer activity of Mzb as a monotherapy in both primary and metastatic TNBCs. We show that Mzb strongly inhibits proteasome functions by blocking all three proteasome activities (CT-L, T-L, and CP-L), induces caspase-3 dependent apoptosis, and inhibits tumor cell proliferation *in vitro* in 2D and 3D cultures. Here, we describe an additional, novel mechanism for the anti-cancer activity of Mzb and demonstrate that Mzb inhibits complex II-dependent mitochondrial respiration leading to reduced oxidative phosphorylation (OXPHOS). We further show that Mzb significantly inhibits primary tumor growth and induces apoptosis in human TNBC cell line xenografts, a murine syngeneic TNBC model, and patient-derived tumor xenograft (PDX) models. Mzb also exerts a synergistic anti-cancer activity in combination with doxorubicin *in vivo.* Our results document that Mzb inhibits TNBC metastasis by inhibiting OXPHOS and reducing the number of circulating tumor cells (CTCs), and also reduces spontaneous lung and brain metastatic burden *in vivo* after resection of the primary TNBC tumors.

## Results

### Marizomib selectively inhibits proliferation of TNBC cells

Since the proteasome pathway is more prominent in TNBCs compared to other subtypes of BC and TNBCs show survival dependency on the proteasome complex (5,10), we evaluated the anti-tumor activity of Mzb in TNBCs and other subtypes of BC. We first investigated the effect of Mzb and Btz on the CT-L, T-L, and CP-L proteasome activities in two TNBC lines, SUM159PT and MDA-MB-231. Mzb at 50 nM inhibited the CT-L, T-L, and CP-L proteasome activities by 50%, which was further reduced to 90% at 100 nM (Fig S1A). In line with a previous study (8), 100 nM Btz only inhibited the CT-L proteasome activity by 50-60% in these cells (Fig S1B). Thus, Mzb inhibits all three proteasome active sites in TNBC cells, exerting an overall stronger proteasome inhibitory effect in TNBCs.

Mzb inhibits tumor growth in multiple cancers including myeloma, glioblastoma, leukemia, colorectal, and pancreatic cancer (13). We therefore evaluated the growth inhibitory effect of Mzb using a panel of TNBC, non-TNBC, and non-malignant mammary epithelial cell lines. Mzb selectively reduced proliferation of TNBC cells in a concentration-dependent manner (Fig 1A), without much effect on non-TNBC and non-malignant mammary epithelial cells

**Fig 1:**
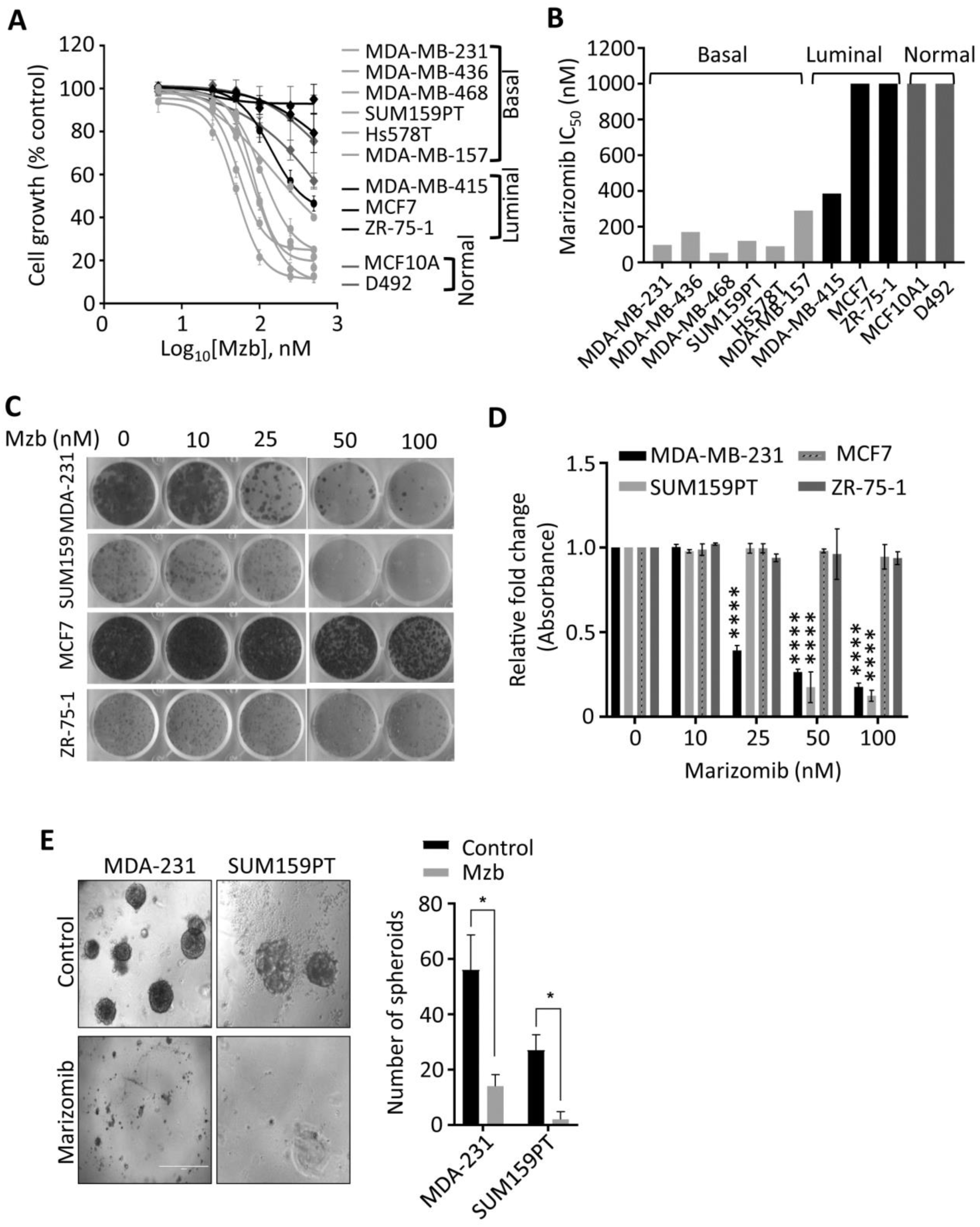
Marizomib selectively inhibits proliferation of TNBC cells in 2D and 3D culture. **(A)** A panel of TNBC (basal), luminal, and non-malignant mammary epithelial cells were treated with Mzb (0-500 nM) and cell proliferation was analyzed after 6 days using the MTS assay. The dose response curve was generated by calculating cell viability relative to the DMSO-treated control. One-way ANOVA and Tukey’s post-test were employed (n=3). **(B)** The IC50 values of Mzb in TNBC (basal), luminal, and non-malignant cells. **(C, D)** Representative images of the colony-forming capacity of TNBC lines (SUM159PT and MDA-MB-231) and luminal BC lines (MCF7 and ZR-75-1) following Mzb treatment (0-100 nM) at 14 days analyzed using crystal violet staining (C). Quantification of the colonies formed in all cell lines following Mzb treatment measured by reading crystal violet absorbance (D). One-way ANOVA and Tukey’s post-test were employed (n=3). **(E)** Left panel, representative images of SUM159PT and MDA-MB-231 spheroids grown on Matrigel for 14 days treated with 100 nM Mzb. Right panel, quantification for a number of tumor spheroids treated with 100 nM Mzb as analyzed by counting the number of spheroids under phase-contrast microscope. The paired Student’s *t* test was performed (n=3).

(MCF10A and D492) (Fig 1A). The IC50 values for Mzb were less than 150 nM in TNBC lines and greater than 1 µM in non-TNBC lines (Fig 1B). We next analyzed the effect of Mzb on the clonogenic activity of TNBC and non-TNBC cells using a long-term colony-forming assay. Mzb significantly reduced the number of colonies for TNBC lines (SUM159PT and MDA-MB-231) but not for non-TNBC lines (MCF7 and ZR-75-1) (Fig 1C), suggesting that Mzb selectively inhibits TNBC cell growth.

We next examined whether Mzb can inhibit tumor growth in a 3D culture setting, which simulates the *in vivo* physiological environment of tumor cells and is used for enriching tumor-initiating cells. We used SUM159PT and MDA-MB-231 TNBC cells that form spheroids under non-adherent conditions. Mzb (100 nM) significantly reduced the size and number of MDA-MB-231 spheroids compared to untreated spheroids (Fig 1D). Interestingly, Mzb completely eliminated SUM159PT spheroid formation (Fig 1D). Thus, Mzb effectively reduces tumor cell growth in both 2D and 3D cultures.

### Marizomib induces caspase-3 dependent apoptosis in TNBC cells

Proteasome inhibitors including Btz and Mzb induce caspase-dependent apoptosis in leukemia and glioblastoma (14,15). We therefore examined whether Mzb induces caspase-dependent apoptosis in TNBC cells. Mzb increased caspase-3 activity, as measured by the cleavage of caspase-3-specific substrate Ac-DEVD-AMC in TNBC cells but not in luminal and non-malignant breast cells (Fig S2A). Moreover, Mzb-induced increase in caspase-3 activity was rescued by co-treatment with the pan-caspase inhibitor z-VAD-FMC in SUM159PT and MDA-MB-231 (Fig S2B). Furthermore, Mzb treatment resulted in a considerable cleavage of PARP1, a classical marker of apoptosis, in TNBCs cells (SUM159PT, MDA-MB-231, and BT-549) (Fig S2C). We also analyzed the effect of Mzb treatment on the expression of various anti-apoptotic, pro-survival, and pro-apoptotic proteins. Our results showed that Mzb reduced the expression of anti-apoptotic proteins Mcl-1 and Bcl-2, and of the survival protein survivin. Notably, Mzb treatment increased the expression of pro-apoptotic protein Bim (Fig S2D). Hence, our data reveal that Mzb induces caspase-3-dependent apoptosis in TNBC cells.

### Marizomib inhibits mitochondrial respiration and OXPHOS in TNBC cells

Proteasome inhibitors such as Btz affect other targets in cancer cells in addition to their primary target, the proteasome complex. Hence, we investigated possible additional targets of Mzb in TNBC cells that may also contribute to the observed anti-cancer activity. To address this question, SUM159PT cells were treated with 100 nM Mzb for 0 and 9 h (before apoptosis induction) and label-free global proteomic analysis was carried out to identify the proteins or pathways altered by Mzb. Our proteomic analysis revealed that Mzb reduced the levels of 11 proteasome subunits (Fig 2A). Interestingly, metabolism pathway components were also significantly downregulated following a 9 h Mzb treatment (Fig 2A). When we looked at individual metabolic pathways, we observed that 18 proteins of the OXPHOS pathway were downregulated (log_2_FC≤ –0.5) following Mzb treatment (Fig 2B, C). We next analyzed the protein levels of one representative component of each of the mitochondrial respiration complexes including NDUFB8 (complex I), SDHB (complex II), UQCRC2 (complex III), COX-II (complex IV), and ATP5A (complex V) by western blotting. Mzb markedly reduced SDHB and NDUFB8 protein levels (Fig 2D), suggesting that Mzb primarily affects mitochondrial complex I and II.

**Fig 2:**
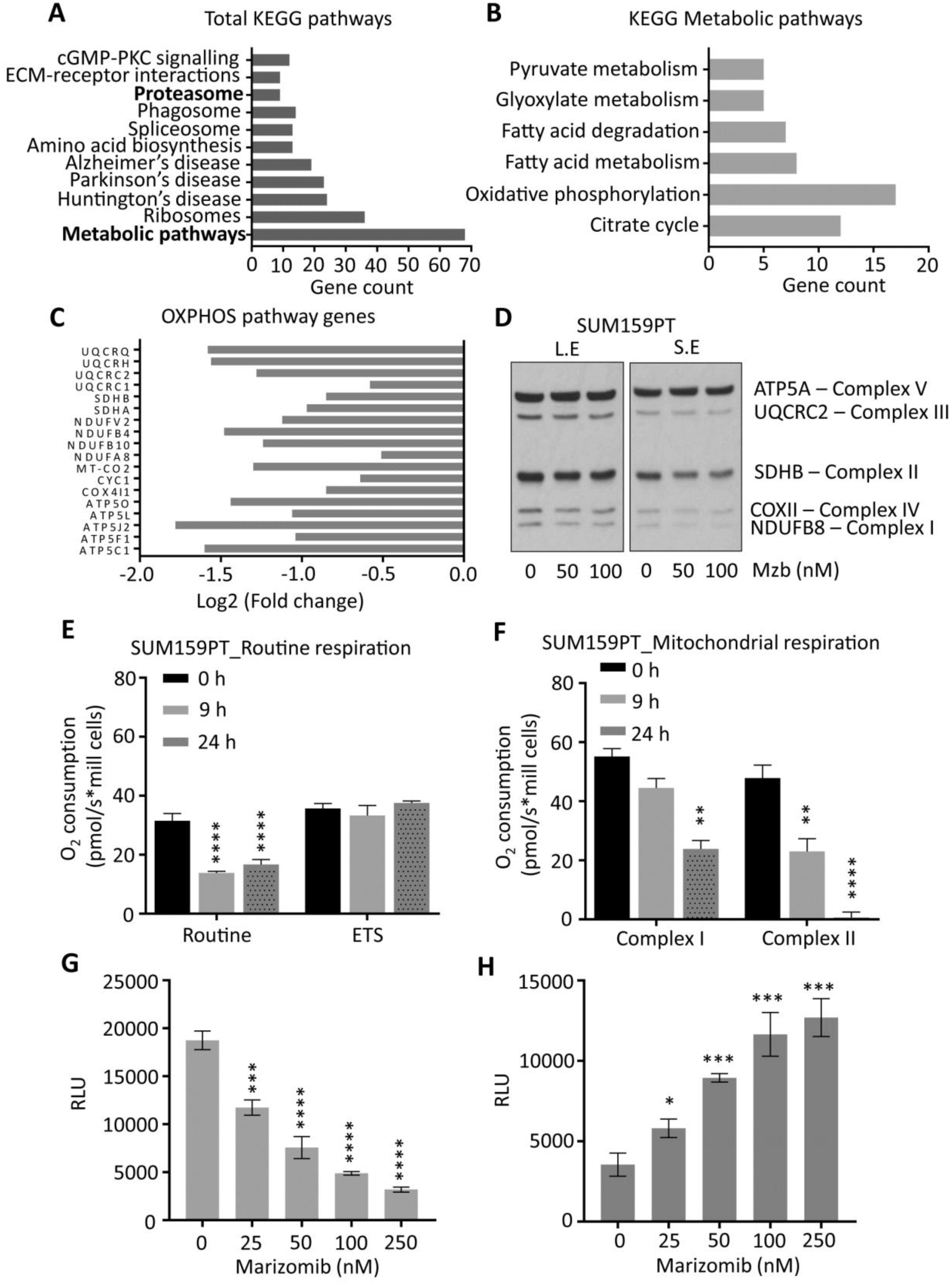
Marizomib inhibits mitochondrial respiration and OXPHOS in TNBC cells. **(A-C)** SUM159PT cells were treated with Mzb (100 nM) for 9 h and label-free global proteomic analysis was carried out. List of total KEGG pathways (A), KEGG metabolic pathways (B), and the OXPHOS pathway proteins (C) downregulated following 9 h of Mzb treatment. **(D)** SUM159PT cells were treated with Mzb (0-100 nM) for 24 h and protein levels of indicated OXPHOS pathway proteins were analyzed by western blotting. Left panel represents a long exposure (L.E) and right panel represents a short exposure (S.E). **(E, F)** SUM159PT cells were treated with Mzb (100 nM) for 9 and 24 h. (E) Oxygen consumption for ROUTINE respiration and FCCP-stimulated uncoupled respiration capacity (ETS) were evaluated on intact cells. (F) Oxygen consumption of the cells was evaluated for Complex I/CII-linked respiration. One-way ANOVA and Tukey’s post-tests were employed (n=4 or 5). **(G, H)** SUM159PT cells were treated with Mzb (0-100 nM) for 24 h and intracellular ATP (G) and ROS (H) levels were analyzed. One-way ANOVA and Tukey’s post-tests were employed (n=3).

Although TNBCs have been shown to rely on glycolysis (16,17), recent studies have shown that the components of the mitochondrial OXPHOS pathway are upregulated in TNBCs, and its suppression by OXPHOS inhibitors reduces TNBC tumor progression (2,18). Hence, we further explored this novel OXPHOS inhibitory activity of Mzb in TNBCs. We evaluated routine respiration and mitochondrial respiration of SUM159PT cells following Mzb treatment using high-resolution respirometry. We found that Mzb significantly reduced routine respiration of the cells by ∼50% after 9- and 24-h treatments, however the maximum respiratory capacity of the cells was unaffected (Fig 2E). In order to better understand the respiration inhibition capability of Mzb, we next examined if Mzb inhibits complex I-dependent or complex II-dependent respiration in SUM159PT cells. While 9 h Mzb treatment had no significant effect on complex I-dependent respiration, it inhibited complex II-dependent respiration by ∼50% (Fig 2F). Moreover, we observed a complete inhibition of complex II-dependent respiration after 24 h Mzb treatment, but only observed a marginal inhibition of complex I-dependent respiration (Fig 2F). Furthermore, Mzb reduced ATP generation (Fig 2G) and increased reactive oxygen species (ROS) formation (Fig 2H) in SUM159PT cells in a dose-dependent manner. Similar to the effects on SUM159PT cells, Mzb reduced routine and complex II-dependent respiration (Fig S3A, B), decreased ATP generation (Fig S3C), and increased ROS levels (Fig S3D) in MDA-MB-231 cells. Hence, Mzb inhibits mitochondrial respiration in TNBC cells in addition to the proteasome complex inhibition.

### Marizomib inhibits OXPHOS in a PGC-1α-dependent manner

We next delineated the mechanism by which Mzb inhibits OXPHOS in TNBC cells. Since PGC-1α, a member of PGC-1 family of transcriptional co-activators, is known to regulate OXPHOS in breast cancer, we examined the effect of Mzb on PGC-1α expression in SUM159PT cells. We found that Mzb (100 nM) markedly reduced PGC-1α protein levels within 9 h treatment (Fig 3A). The reduced levels of PGC-1α are most likely due to reduction in mRNA levels of PGC-1α after Mzb (100 nM) treatment (Fig 3B). Moreover, the mRNA levels of components of complex-I and II OXPHOS genes (SDHA, SDHB, NDUFB8, and NDUFB4) (Fig 3B), downstream targets of PGC-1α, were also reduced upon 9 h Mzb treatment. To further confirm the role of PGC-1α in Mzb-induced OXPHOS inhibition and cell death, we exogenously overexpressed PGC-1α in SUM159PT cells and subsequently treated them with Mzb (100 nM) for 9 h to assess OXPHOS gene expression and for 24 h to assess cell viability. Interestingly, PGC-1α overexpression partially rescued SUM159PT cells from undergoing Mzb-induced cell death (Fig 3C) and also rescued Mzb-induced OXPHOS gene downregulation (Fig 3D). Hence, our data indicates that Mzb exerts its anti-cancer activity in-part via PGC-1α-mediated OXPHOS inhibition.

**Fig 3:**
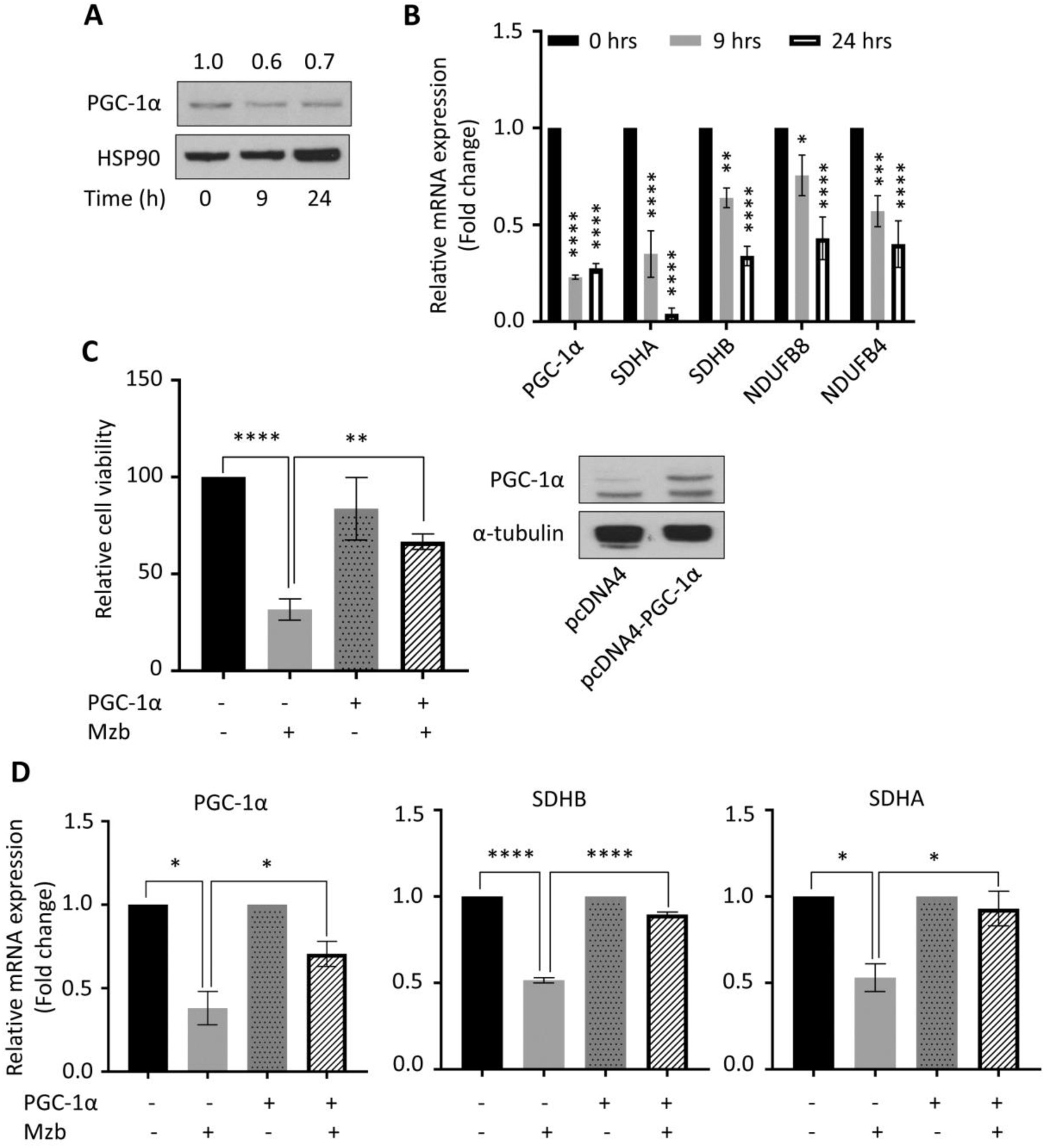
Marizomib inhibits OXPHOS in a PGC-1α-dependent manner. **(A)** SUM159PT cells were treated with 100 nM Mzb for 0, 9, and 24 h, and PGC-1α protein levels were analyzed. HSP90 was used as a loading control. **(B)** SUM159PT cells were treated with 100 nM Mzb for indicated time-points, and mRNA levels of the indicated genes were analyzed by RT-qPCR. One-way ANOVA and Tukey’s post were used (n=3). **(C, D)** SUM159PT cells were transfected with either pcDNA4 vector or pcDNA4-Myc-PGC-1α overexpression plasmid for 24 hours, and subsequently treated with 100 nM Mzb for 9 (for OXPHOS gene expression) and 24 h (for cell viability). (C) Left panel, cell viability was analyzed by Trypan blue exclusion assays. (D) OXPHOS gene expression levels were analysed by RT-qPCR. One-way ANOVA and Tukey’s post-test were performed (n=2). Right panel, representative western blot images showing PGC-1α overexpression.

### Marizomib suppresses TNBC tumor growth *in vivo*

Since FDA-approved peptide-based proteasome inhibitor Cfz failed to suppress tumor growth in a human breast cancer cell line MDA-MB-231-derived xenograft model (5), we evaluated the anti-cancer activity of Mzb in TNBC models *in vivo*. For this purpose, we first used MDA-MB-231 xenografts. Mzb treatment significantly reduced tumor volume (Fig 4A) and tumor weight (Fig 4B) compared to the vehicle-treated group, suggesting the superior anti-cancer activity of Mzb in TNBCs. We then analyzed the anti-tumor activity of Mzb using a TNBC patient-derived tumor xenograft (PDX). Interestingly, Mzb treatment significantly inhibited tumor growth (Fig 4C) and reduced tumor weight (Fig 4D) in our PDX model. More specifically, four out of six PDX tumors regressed following Mzb treatment (Fig 4C).

**Fig 4:**
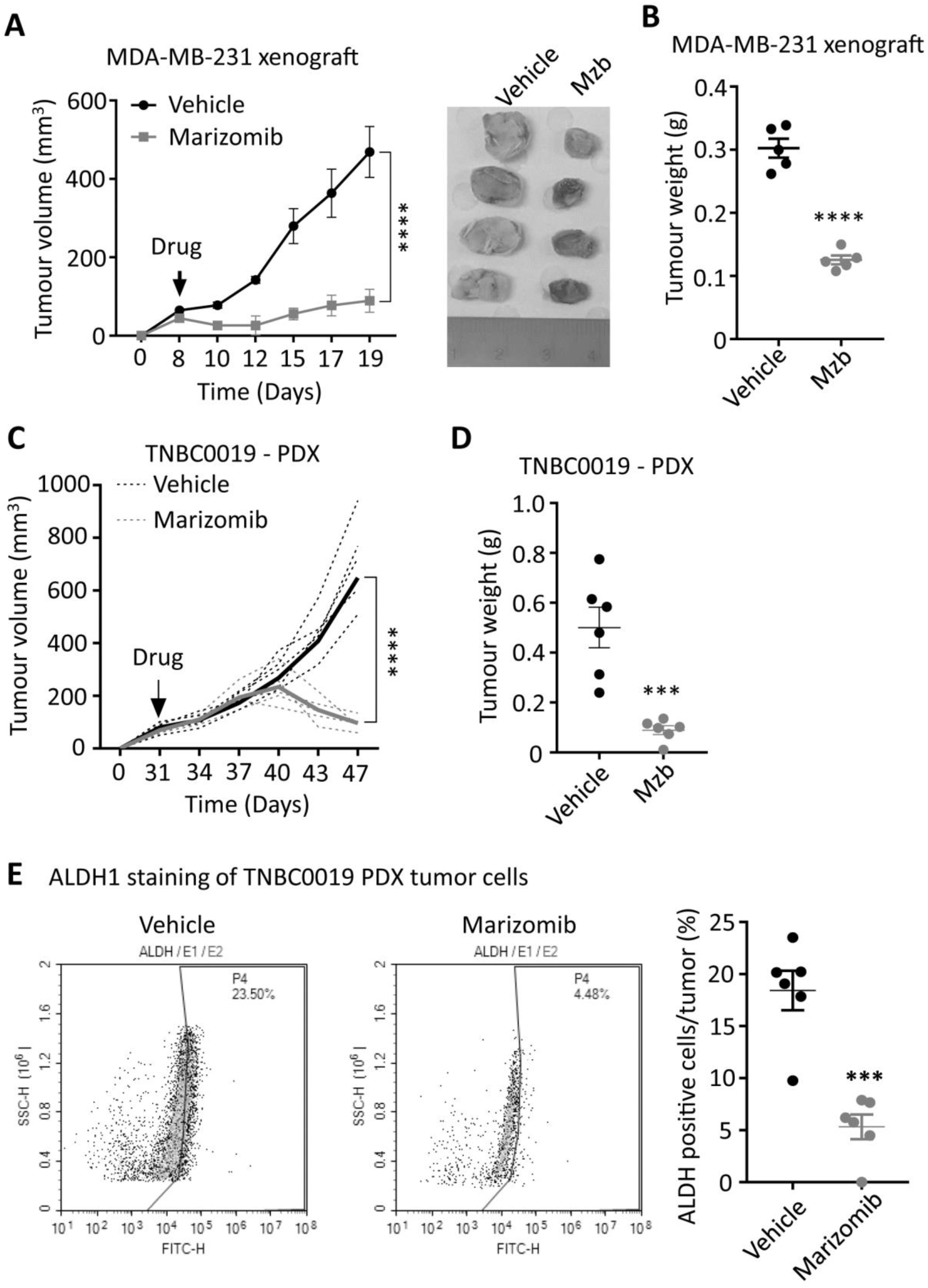
Marizomib suppresses TNBC tumor growth *in vivo* in MDA-MB-231 xenograft and patient-derived tumor xenograft. **(A, B**) MDA-MB-231 tumor growth following treatment with the vehicle or Mzb (0.15 mg/kg, twice/week, IP) for two weeks. The mean tumor size of each treatment group (A) is presented. Tumor weight (B) measured at the end of the two weeks treatment is presented (n=6 mice/group). **(C, D)** TNBC PDX tumor growth following vehicle or Mzb administration (0.15 mg/kg, twice/week, IP) 2-week treatment. The growth of individual PDX tumors (dotted line) and their mean tumor size (solid line) is presented (C). Tumor weight (D) was measured at the end of the (n=6 mice/group). Two-way ANOVA and Sidak’s post-test were used for tumor growth analysis. Unpaired “*t*” test was used for tumor weight analysis. **(E)** PDX tumors treated with vehicle or marizomib for 2 weeks were stained with the antibody against human ALDH1, and BCSC populations were analyzed by flow cytometry (n=6, mice/group). Representative images of contour plot of ALDH1^+^ (left panel). Quantification of ALDH1^+^ subpopulation of cells is presented in right panel.

One of the reasons for the lack of an effective therapy for TNBC patients is the inability of the current therapies to effectively eradicate tumor-initiating cells, also known as breast cancer stem cells (BCSCs). BCSCs are inherently resistant to the standard-of-care neoadjuvant chemotherapies and radiotherapy, resulting in therapy resistance and metastasis (19-22). Since Mzb treatment significantly regressed patient-derived tumors *in vivo* (Fig 4C), we analyzed BCSC population in the tumor tissues resected at the end of a two-week treatment. The resected patient-derived control tumors (vehicle-treated) were enriched with an ALDH1^+^ population. Mzb treatment significantly reduced the percentage of ALDH1^+^ BCSCs from approximately 23% in control tumors to 4-5% (Fig 4E), suggesting that Mzb does not only impact the bulk-tumor population but also impacts BCSCs, and therefore regressed patient-derived tumors *in vivo*.

We next tested the anti-cancer activity of Mzb using a fully immunocompetent murine pre-clinical syngeneic model. 4T1.2 is highly aggressive murine TNBC cell line that aptly recapitulates human TNBC tumor phenotypes after orthotopic mammary fat pad implantation in immunocompetent Balb/c mice. Our results show that Mzb significantly reduced 4T1.2 tumor growth (Fig 5A) and tumor weight (Fig 5B) compared to the vehicle-treated group. We also analyzed the percentage of apoptotic cells in vehicle- and Mzb-treated tumors by ApopTag staining and observed an increased percentage of ApopTag-positive cells in Mzb-treated 4T1.2 tumor tissues (Fig 5C), suggesting that Mzb induces apoptosis *in vivo*. Since 4T1.2 model is invasive and forms spontaneous lung metastasis, we examined whether Mzb was able to inhibit lung metastasis in the 4T1.2 model. Hematoxylin and eosin (H&E) staining of the lungs isolated from vehicle- and Mzb-treated mice after two weeks of treatment showed a significant reduction in the number of lung metastatic nodules in Mzb-treated mice compared to vehicle-treated mice (Fig 5D). We then examined the effect of Mzb on the expression of various epithelial-to-mesenchymal (EMT) markers and cell migration *in vitro*. Mzb treatment (100 nM) reduced mRNA levels of multiple EMT markers including ZEB1, Vimentin, and Slug in 4T1.2, SUM159PT, and MDA-MB-231 cells (Fig S4A-C). Mzb also reduced SUM159PT and MDA-MB-231 cell migration *in vitro* (Fig S4D). Thus, our data indicate that Mzb reduces TNBC tumor growth and inhibits metastases.

**Fig 5:**
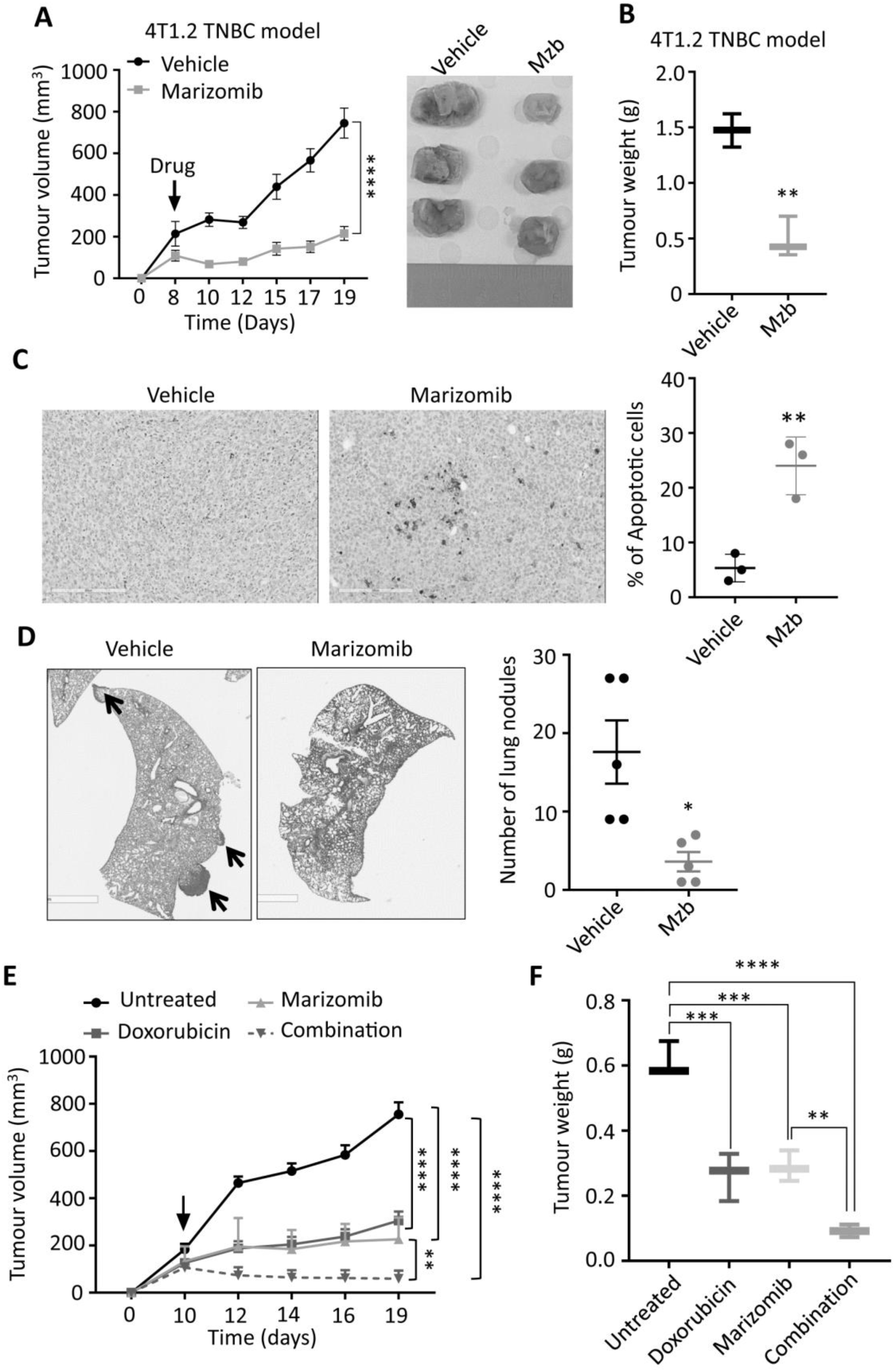
Marizomib inhibit tumor growth in syngeneic 4T1.2 murine model of TNBC *in vivo*. **(A, B)** Murine 4T1.2 syngeneic TNBC tumor growth following treatment with vehicle or Mzb (0.15 mg/kg, twice/week, IP) for two weeks is shown. The mean tumor volume (A) of each treatment group and tumor weight (B) from each mouse is presented (n=6 mice/group). **(C)** Left, representative image of ApopTag staining of primary 4T1.2 tumors treated with the vehicle or Mzb for 2-weeks. Right, quantification for percentage of ApopTag-positive cells in primary 4T1.2 tumors following Mzb treatment. Unpaired “*t*” test was performed (n=3). **(D)** Left panel, representative images of lung metastasis in 4T1.2 tumor model treated with the vehicle or Mzb for two weeks. The metastatic nodules were stained with H&E. Two-way ANOVA followed by Sidak’s post-test were employed for tumor growth analysis and the paired student “*t*” test was employed for tumor weight analysis and lung metastasis analysis (n=5). **(E, F)** Murine 4T1.2 syngeneic TNBC tumor growth following treatment with the vehicle, Mzb (0.075 mg/kg), doxorubicin (5 mg/kg), and combination for two weeks. The mean tumor volume of each treatment group and tumor weight (F) from each mouse is presented (n=6 mice/group). Two-way ANOVA followed by Sidak’s post-test for tumor growth analysis and one-way ANOVA followed by Tukey’s post-tests for tumor weight analysis were performed.

We next evaluated the anti-cancer activity of Mzb in combination with a standard-of-care chemotherapy, doxorubicin, in a syngeneic 4T1.2 model of TNBC. Mice were treated with half-maximum tolerated dose (MTD) of Mzb (0.075 mg/kg) or doxorubicin (5 mg/kg) alone or in combination. Both Mzb and doxorubicin when used as monotherapy significantly delayed tumor growth compared to untreated tumors. Notably, we observed that combination of Mzb and doxorubicin significantly inhibited 4T1.2 tumor growth (Fig 5E, F) compared to Mzb and doxorubicin monotherapy. The combination was well-tolerated as no significant weight loss was observed. Thus, our data suggests that combining Mzb with doxorubicin could serve as a better therapeutic strategy for improved clinical outcomes in TNBC patients.

### Marizomib reduces spontaneous metastasis in murine TNBC model by inhibiting EMT and circulating tumor cells

Although Mzb inhibited lung metastasis in 4T1.2 tumor model (Fig 4D), it is possible that the effect on metastasis in this model is due to suppression of primary tumor growth (Fig 5A). To overcome the potential confounding effects on primary tumor growth, we next investigated the effect of Mzb on spontaneous or established lung and brain metastasis using a more aggressive syngeneic murine 4T1BR4 TNBC model. The 4T1BR4 cells are derived from the parental 4T1 cells with approximately 20% uptake rate in the brain and 60% uptake rate in the lungs (23). Following the engraftment of 4T1BR4 cell and once the tumor size reached ∼250-300 mm^3^, we surgically resected the primary tumors. Two days after resection, mice were treated with the vehicle or Mzb (0.15 mg/kg) for two weeks (Fig 6A). At the end of the treatment, lung and brain metastasis were analyzed. Our data revealed that Mzb dramatically reduced the number of micro and macro lung metastatic nodules in 4T1BR4 tumor model (Fig 6B). Similarly, Mzb reduced the number of micro brain metastatic nodules (Fig 6C). We were unable to assess the effect of Mzb on macro-brain metastasis as these are not evident in the 4T1BR4 model. Our data convincingly show that Mzb exerts potent cytotoxicity on metastatic TNBCs by reducing the overall lung metastasis burden and micro metastasis in the brain.

**Fig 6:**
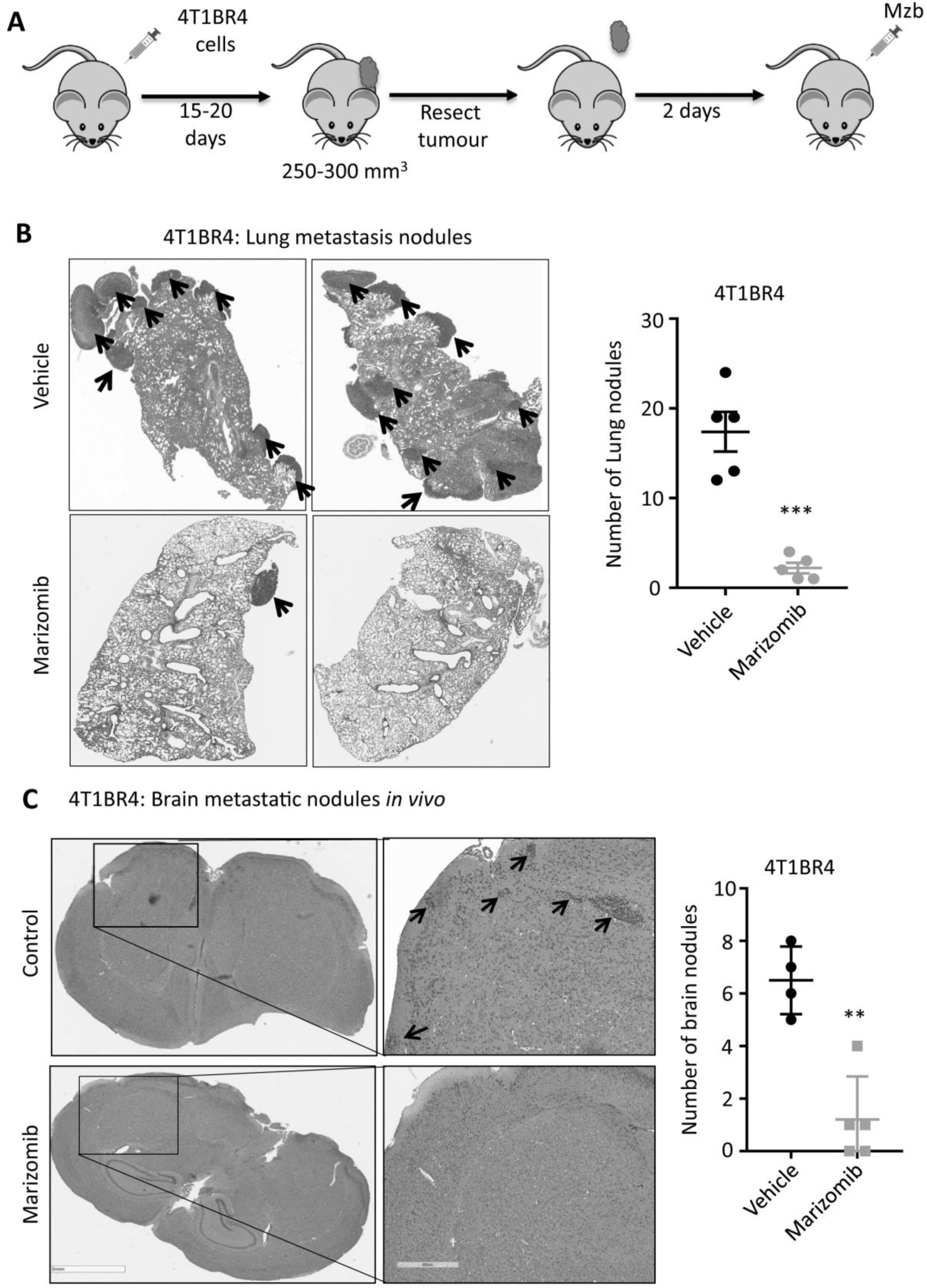
Marizomib inhibits spontaneous lung and brain metastasis *in vivo.* **(A)** Schematic representation of the 4T1BR4 tumor resection model to study the effect of marizomib on established lung and brain metastasis. **(B)** Representative images of metastatic 4T1BR4 tumors in the lungs (left panel) after resecting primary tumors and following the vehicle or Mzb treatment. The metastatic nodules were stained with H&E staining (n=5 mice/group). Number of micro and macro metastatic 4T1BR4 nodules in the lungs (right panel) are shown. **(C)** Representative images of metastatic 4T1BR4 tumors in the brain (left panel) after resecting primary tumors and following vehicle or Mzb treatment. The metastatic nodules were stained with H&E staining (n=5 mice/group). Number of micro metastatic 4T1BR4 nodules in the brain (right panel) are shown (n=5 mice/group). Unpaired student “*t*” test was employed.

Mzb has been shown to reduce prostate tumor metastasis by suppressing the expression of EMT markers via NF-κβ inhibition (24). In our study we also found that Mzb-induced a significant downregulation of the key regulator of cell metabolism PGC-1α, which has been shown to be upregulated in metastatic breast cancer cells and to promote breast cancer metastasis by stimulating OXPHOS, increasing cell motility and invasive properties without impacting the expression of EMT-related genes (25), suggesting that Mzb catalyzed suppression of PGC-1α levels is not mutually exclusive to suppression of EMT markers. We examined the effect of Mzb on the expression of PGC-1α and EMT markers in 4T1BR4 primary tumors treated with the vehicle or Mzb for one week. This treatment significantly reduced the number of lung metastatic nodules in 4T1BR4 model *in vivo* (Fig 7A). Notably, Mzb significantly reduced the mRNA levels of PGC-1α and EMT genes including ZEB1, vimentin, and slug (Fig 7B) in primary tumors. Moreover, PGC-1α and EMT genes are known to regulate the number of CTCs, and promote breast cancer metastasis (26). Next, we examined the effect of Mzb on CTCs. We isolated CTCs from the blood of both vehicle- and Mzb-treated mice presented in Fig 7A. Interestingly, Mzb dramatically reduced CTCs in the 4T1BR model *in vivo* (Fig 7C). Hence, our data suggested that Mzb may inhibit TNBC lung metastasis via repressing PGC-1α and consequent OXPHOS inhibition in addition to suppressing expression of EMT markers.

**Fig 7:**
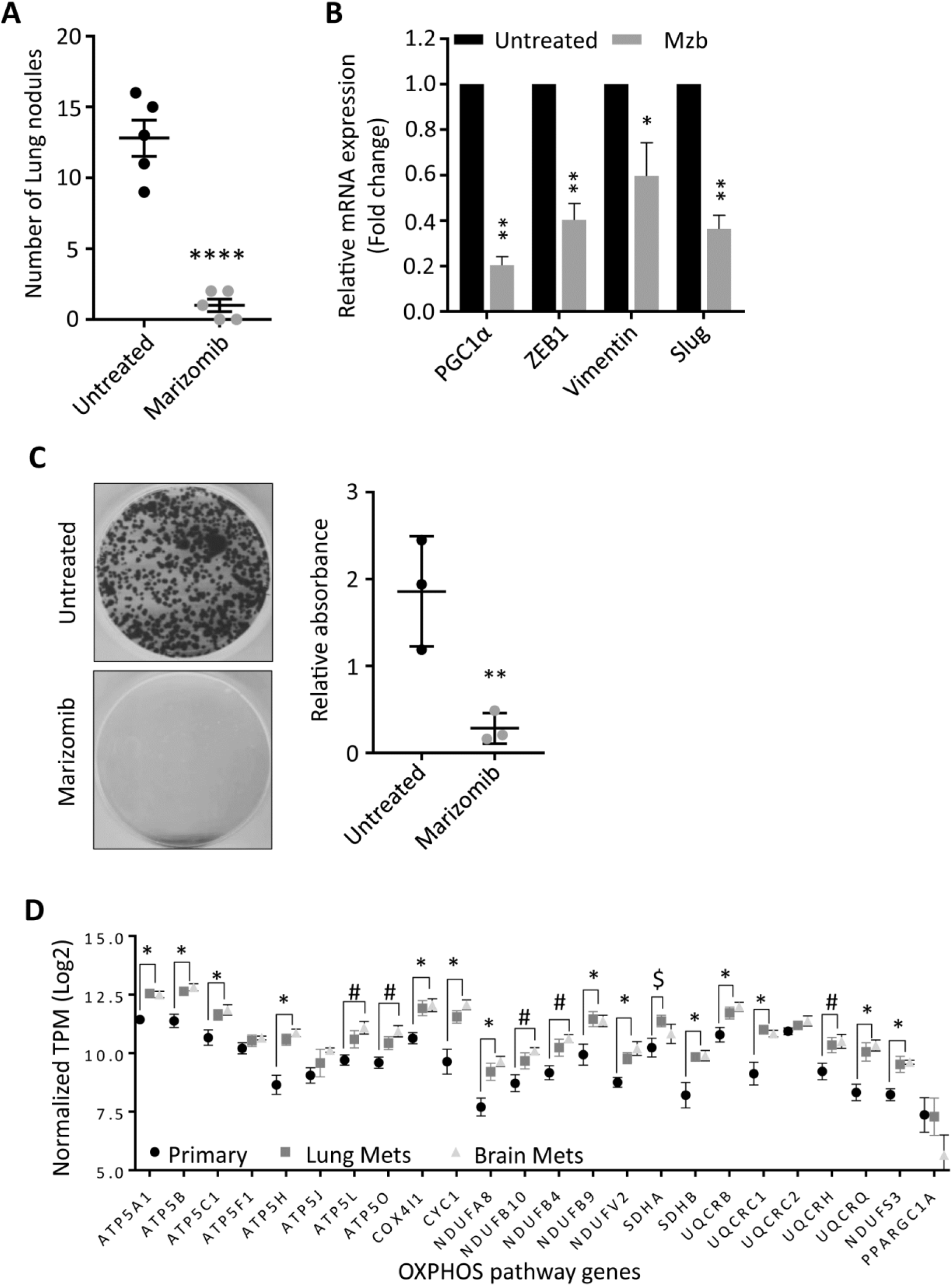
Marizomib reduces metastasis by inhibiting EMT and circulating tumor cells. **(A)** The number of lung metastatic nodules stained with H&E staining in primary 4T1BR4 tumors treated with the vehicle or Mzb (0.15 mg/kg) for one week. The unpaired “*t*” test was performed. **(B)** Expression levels of PGC-1α, ZEB1, Vimentin, and Slug mRNA in vehicle- and Mzb-treated primary 4T1BR4 tumors (n=3) were analyzed by RT-qPCR. **(C)** Left panel, representative images for circulating tumor cells isolated from the blood of vehicle- or Mzb-treated mice implanted with 4T1BR4 tumors. Right panel, quantification of circulating tumor cells colonies stained with crystal violet (n=3). The unpaired “*t*” test was performed. **(D)** Expression of an indicated OXPHOS genes in matched primary, lung and brain metastatic breast cancer patients (n=12). Unpaired “*t*” test was performed. * (primary Vs lung and brain metastasis), (primary Vs lung metastasis), and $ (primary Vs brain metastasis).

Due to an established role of OXPHOS in TNBC metastasis in pre-clinical models (26), we next examined the expression of candidate OXPHOS genes in the breast cancer patient dataset GSE110590 (27). We analyzed the absolute expression values of RNA derived from RNA sequence normalized counts (RSEM) from matched primary breast cancer, brain metastases and lung metastases (n=6 for lung metastases, n=5 for brain metastases). Interestingly, our results show that the expression of OXPHOS genes was significantly higher in lung and brain metastatic patients compared to their matched primary tumors (Fig 7D). These data suggest that OXPHOS pathway is enriched in metastatic patients and may potentially drive metastasis in TNBC patients, and therefore Mzb could provide an effective therapy option for metastatic TNBC patients or prevent formation of metastatic tumors in patients.

### Marizomib exerts a synergistic anti-cancer activity with glycolysis inhibitor

In response to OXPHOS inhibition, cancer cells switch to glycolysis to meet their high energetic demand and survival (28). We therefore examined if Mzb upregulates glycolysis in TNBC cells when OXPHOS is inhibited. We analyzed our proteomic data and searched for the pathways upregulated in response to Mzb in SUM159PT cells and found a significant upregulation of two metabolic pathways, glycolysis and carbon metabolism (Fig 8A). Since glycolysis has been previously shown to compensate for the loss of the OXPHOS pathway (28), we focused on it for further analysis. We identified a significant upregulation of 9 components of the glycolysis pathway including LDHA, HK1, PGK1, and GAPDH following Mzb treatment (Fig 8B). This suggests that upregulated glycolysis may compensate for the loss of OXPHOS to fuel the energetic demands of TNBC cells. Furthermore, Mzb treatment (100 nM) increased intracellular lactate levels in SUM159PT cells, which was reduced by STF-31 (0.5 µM), a known GLUT1 inhibitor (Fig 8C). Hence, TNBC cells upregulate glycolysis to compensate for the loss of OXPHOS following Mzb treatment.

**Fig 8:**
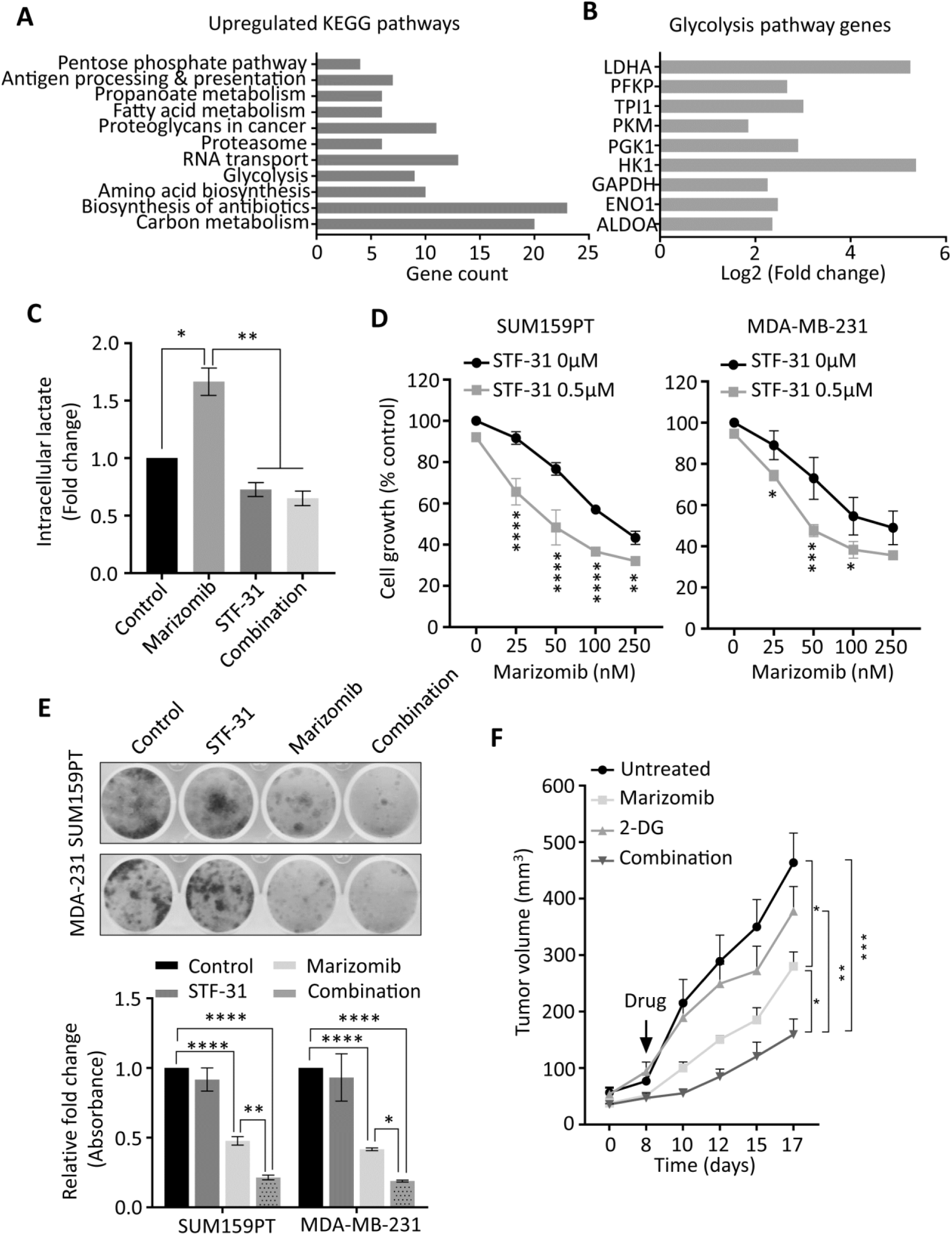
Marizomib exerts a synergistic anti-cancer activity with glycolysis inhibitor. **(A, B)** List of KEGG pathways (A) and glycolysis pathway proteins (B) upregulated in SUM159PT cells following 9 h Mzb (100 nM) treatment. **(C)** SUM159PT cells were treated with Mzb (100 nM) and STF-31 (0.5 µM), alone or in combination for 16 h. Intracellular lactate levels were analyzed by the Lactate-Glo assay kit. One-way ANOVA followed by Tukey’s post-tests were employed (n=3). **(D)** SUM159PT and MDA-MB-231 cells were treated with Mzb (0-250nM) with or without STF-31 (0.5 µM) for 4 days, and cell viability was analyzed by the MTS assays. Two-way ANOVA and Sidak’s post-test were employed (n=3). **(E)** Representative images of colony-forming capacity of SUM159PT and MDA-MB-231 cells following the treatment with Mzb (100 nM) and STF-31 (0.5 µM), both alone and in combination, at 14 days analyzed using crystal violet staining (upper panel). Quantification of the colonies formed in both cell lines measured by reading crystal violet absorbance (lower panel). One-way ANOVA and Tukey’s post-test were employed (n=3). **(F)** Murine 4T1.2 syngeneic TNBC tumor growth following the treatment with vehicle, Mzb (0.075 mg/kg), 2-DG (400 mg/kg), and combination for two weeks. The mean tumor volume of each treatment group is presented (n=6 mice/group). Two-way ANOVA followed by Sidak’s post-test were employed.

We next examined the effect of glycolysis inhibition in combination with Mzb on TNBC cell viability. We found that Mzb exerted a synergistic growth inhibitory effect in combination with STF-31 in SUM159PT and MDA-MB-231 cells (Fig 8D). We then evaluated the effect of Mzb-STF-31 combination on the clonogenic activity of SUM159PT and MDA-MB-231 cells. Our data showed that Mzb-STF-31 combination therapy significantly reduced the number of colonies for both cell lines (Fig 8E).

We next investigated the effect of Mzb in combination with a glycolysis inhibitor *in vivo* using 4T1.2 model. For this study, we used 2-deoxy D-glucose (2-DG), since STF-31 is not suitable for *in vivo* work. Our results show that Mzb in combination with 2-DG significantly inhibited 4T1.2 tumor growth, as assessed based on tumor volume and weight, compared to Mzb or 2-DG monotherapy (Fig 8F, G). This combination was well tolerated as indicated by the absence of weight loss. Notably, although Mzb monotherapy (0.075 mg/kg) significantly inhibited tumor growth, 2-DG alone had no inhibitory effect on 4T1.2 tumors. Collectively, our data indicate that upon OXPHOS inhibition, TNBC cells switch to glycolysis, and combined inhibition of glycolysis with OXPHOS appears to be rather efficient.

## Discussion

In this study, we show for the first time that Mzb strongly inhibits TNBC primary tumor growth and reduces its metastasis in the lungs and micro-metastasis in the brain. Further, we document that Mzb, in addition to targeting the proteasome complex, inhibits complex II-dependent mitochondrial respiration underlying its cytotoxic activity in both primary and metastatic TNBCs. Finally, we found that Mzb exerts a synergistic anti-cancer activity with doxorubicin, a standard-of-care therapy for TNBC patients, as well as with the glycolysis inhibitor 2-DG.

Although TNBC cells show survival dependency on the proteasome genes and its catalytic activity (2), FDA-approved peptide-based proteasome inhibitors, Btz and Cfz, provided a limited anti-cancer efficacy in pre-clinical models of TNBC and also in clinical trials in breast cancer patients(29,30). Initially, it was believed that poor tumor penetration was a reason for the lack of clinical activity of Btz in solid tumors. However, pharmacodynamics data from two clinical trials revealed that Btz-induced proteasome inhibition is comparable in solid tumors and hematological malignancies (31,32). Both Btz and Cfz inhibit the CT-L proteasome activity without any significant effect on T-L and CP-L activities (12). Inhibition of CT-L by Btz or Cfz leads to a compensatory activation of T-L and CP-L activities, resulting in Btz/Cfz resistance in patients (10,12). Consistent with this, co-inhibition of multiple catalytic sites could overcome Btz resistance in pre-clinical models of TNBCs (10). Hence, a more robust proteasome inhibition by blocking its multiple catalytic sites may represent a more effective therapeutic approach in solid tumors including TNBC.

In clinical settings, Btz and Cfz only inhibit the CT-L activity by 65% and 74%, respectively (33,34), without any effect on T-L and CP-L activities, in whole blood of patients. In contrast, Mzb completely blocked CT-L, T-L, and CP-L proteasome activities in whole blood of myeloma and glioblastoma patients (12). In line with these studies, we show that Mzb at much lower concentrations inhibited the CT-L, T-L, and CP-L activities in TNBC cells (Fig S1). In contrast, Btz only inhibited the CT-L activity at much higher concentration in TNBC cells (Fig S1). Hence, our findings indicate that Mzb can effectively inhibit the proteasome function in TNBC patients.

In this study, we show for the first time an additional mechanism for the anti-cancer activity of Mzb in TNBC cells. Our data demonstrate that Mzb downregulates OXPHOS proteins in-part through the downregulation of PGC-1α, inhibits complex II-dependent mitochondrial respiration, induces intracellular ROS formation, and inhibits mitochondrial ATP generation (Fig 2). Interestingly, TNBC show survival dependency on the genes of proteasome and OXPHOS pathway (2). Although TNBCs have been shown to rely largely on glycolysis for meeting their high metabolic demands and for their survival, studies have shown that TNBCs also utilize the OXPHOS system for survival (2,18,35). Hence, targeting OXPHOS has emerged as a novel therapeutic strategy to combat TNBCs. In line with this notion, our data indicate that Mzb, by simultaneously inhibiting the proteasome activity and OXPHOS, provides an effective therapeutic option for TNBC patients.

Marizomib is currently being tested in several hematological malignancies and solid tumors both as a monotherapy, and in combination with standard-of-care chemotherapy. Mzb monotherapy reduced colorectal cancer growth and sensitized the tumors to 5-fluorouracil and oxaliplatin (36). Mzb also inhibited tumor growth in pancreatic cancer models (37). In addition, Mzb exerted a significant anti-tumor activity in glioblastoma xenograft models, both as a single agent and in combination with HDAC inhibitors and temozolomide (15,38). In line with these studies, our results show that Mzb significantly reduces tumor growth and induces apoptosis in multiple TNBC pre-clinical models including the human cell line MDA-MB-231 xenografts, the murine 4T1.2 syngeneic model, and in PDX models (Fig 4 & 5). In contrast to Mzb, Cfz as a single agent showed no tumor suppressive effect in an MDA-MB-231 xenograft model (5). Mzb, when used at the half-MTD level, exerted a significant anti-tumor activity in combination with doxorubicin, a standard-of-care chemotherapy for TNBC patients, in our syngeneic murine 4T1.2 model (Fig 5). Thus, our data convincingly demonstrate that Mzb exerts a superior anti-cancer activity in TNBC xenograft models and, therefore, it is likely to show an activity in TNBC patients both as a monotherapy and a combinatorial therapy.

Marizomib reduced tumor cell proliferation in 2D culture and 3D tumor spheroid culture (Fig 1). Tumor-initiating cells (TICs) play a significant role in driving tumor progression, therapy resistance, and metastasis. Usually, standard-of-care chemotherapies can effectively eliminate non-TICs but lack effectiveness against TICs, which then leads to chemoresistance and recurrence of the disease. It is therefore of paramount importance to eradicate TIC populations to achieve a considerable clinical response in patients. In breast cancer, targeting BCSCs has been shown to be an effective treatment approach as evidenced in HER2^+^ BC, where combination therapy targeting the IL6 receptor and HER2 effectively abrogated tumor growth and metastasis by eliminating BCSCs (39). In line with these observations, our study indicates that Mzb significantly targets and reduces ALDH1^+^ BCSCs in TNBC PDX model (Fig 4). Increasing evidence suggests that mitochondrial OXPHOS is the preferred form of energy production in TICs. Many studies have shown that TICs, such as CD133^+^ cells of glioblastoma and pancreatic ductal adenocarcinoma, ROS^low^ quiescent leukemia stem cells, lung and breast cancer stem cells as well as mesothelioma-initiating cells show upregulated OXPHOS signature and respiration, relying preferentially on the OXPHOS phenotype for their high metabolic demands (40-44). In addition, ALDH1^+^ BCSCs have been shown to have upregulated OXPHOS pathway and rely on mitochondrial respiration for survival (45). Based on this information and on our data showing that Mzb inhibits OXPHOS in TNBC cells, also targets ALDH1^+^ BCSCs in PDX tumors, we propose that Mzb may reduce the number of BCSCs or TICs via OXPHOS inhibition in TNBCs. However, the detailed mechanism for the effect Mzb on mitochondrial respiration in TNBC-initiating cells or ALDH1^+^ BCSCs needs further investigation.

Our study revealed compelling novel data that Mzb not only suppresses primary TNBC tumor, but also reduces lung and brain metastases in our TNBC syngeneic model. TNBC is associated with an early risk of recurrence, high incidence of lung and brain metastasis, and overall poor survival outcomes (46,47). It is estimated that more than 50% of patients diagnosed with primary TNBCs will develop the metastatic disease with a survival rate of <5-years (48). Furthermore, nearly 46% of women with metastatic TNBCs develop brain metastases, which is associated with a median survival time of 4.9 months (46). The greatest clinical hurdle is how to efficiently treat metastatic TNBCs, especially when accompanied by brain metastases, since there is a lack of drugs capable of crossing the blood-brain barrier and exerting a potent cytotoxic activity. Mzb crosses the blood-brain barrier in a glioblastoma xenograft model and exerts cytotoxic effect on tumor cells (49). Consistent with this study, Mzb treatment significantly reduced the micro-metastatic spread of TNBCs to the brain when tested in a murine 4T1BR4 syngeneic model following primary tumour resection. This is significant, since current imaging modalities lack the capability of capturing micro-metastases. Therefore, a drug that reduces micro-metastasis formation will be important in shaping the prognosis of patients with TNBC. These data also imply that Mzb can cross the blood-brain barrier, reach the TNBC brain metastatic tumor, and exert its cytotoxic effect. Taken together, our data provide evidence that Mzb will not only reduce the primary tumor growth but will also prevent metastatic spread from primary TNBC tumor.

Our findings also show that Mzb significantly reduces lung metastasis in the aggressive and highly metastatic 4T1BR4 syngeneic model of TNBC (Fig 7). Metastasis is driven by invading cancer cells represented by CTCs. CTCs rely on OXPHOS for their energy supply during their transit to distal organs. Further, we have previously shown that CTCs have high respiration than cells derived from corresponding primary tumor, using a 4T1 cancer model (50). Increase in the OXPHOS function linked to the metastatic disease was reported to be dependent on PGC-1α (26). PGC-1a downregulation was shown to reduce lung metastasis by suppressing the expression of various OXPHOS genes without any significant effect on EMT genes (26), suggesting that in breast cancer settings PGC-1α may drive metastasis independent of EMT. Molecular analysis of primary 4T1BR4 tumors revealed that Mzb reduced PGC-1α mRNA levels and also significantly reduced the number of CTCs in 4T1BR4 model (Fig 7). These data suggest that Mzb inhibits PGC-1α-stimulated OXPHOS in TNBC cells which may in-part account for the reduction in invasive or circulating tumor cells and reduced metastasis seen after treatment. However, further studies are required to specifically evaluate mitochondrial respiration and OXPHOS pathway components in CTCs isolated from Mzb-treated mice. Since the number of CTCs in Mzb-treated mice are rather low, we were unable to assess their mitochondrial respiration. Furthermore, as shown previously in prostate cancer, Mzb also reduced levels of various EMT markers including ZEB1, Vimentin, and Slug in TNBC cells, suggesting that Mzb may inhibit the TNBC metastatic disease via multiple mechanisms.

Marizomib is currently being tested in multiple phase I, II, and III clinical trials for refractory multiple myeloma, leukemia, lymphoma, glioblastoma, and malignant glioma (NCT03345095, NCT02330562, NCT02103335, NCT02903069), both as a single agent and in combination therapies. The results from these clinical trials show that Mzb (both as a monotherapy and in combination therapy) is well-tolerated, and demonstrate its promising activity in multiple cancer types (51). Our pre-clinical data in this study generated using a range of *in vitro* and *in vivo* TNBC models provide a strong rationale to translate Mzb into a clinical trial in combination with standard-of-care chemotherapy for primary and metastatic TNBC patients.

## Materials and Methods

### Cell lines and reagents

All breast cancer cell lines and non-malignant mammary epithelial cell line (MCF10A) were obtained from the American Type Culture Collection (ATCC). The 4T1.2 and 4T1BR4 cell lines were kindly provided by Dr Norman Pouliot, Olivia Newton-John Cancer Research Institute, Australia. All breast cancer cell lines were cultured in DMEM media supplemented with 10% fetal bovine serum (FBS). D492 mammary epithelial cells were provided by Dr Thorarinn Gudjonsson (The Panum Institute, Denmark) and were maintained as described previously (52). All cell lines were tested for Mycoplasma infection and authenticated using short tandem repeat (STR) profiling by scientific services at QIMR Berghofer Medical Research Institute. Marizomib was purchased from Cayman Chemicals (Cat #: 10007311).

### Mitochondrial Respiration

Mitochondrial respiration was assessed as described previously (53). Briefly, SUM159PT or MDA-MB-231 were grown to 50-70% confluence, treated with Mzb (100 nM) for 9 h and 24 h, respectively. Cells were harvested and suspended in the DMEM medium without serum. Oxygen consumption for ROUTINE respiration, oligomycin-inhibited LEAK respiration, FCCP-stimulated uncoupled respiration capacity (ETS) and rotenone/antimycin-inhibited residual respiration (ROX) were evaluated using the Oxygraph-2k instrument (Oroboros) on intact cells. The results were normalized to the cell numbers.

For CI- and CII-dependent respiration, cells were harvested and suspended in the mitochondrial respiration medium MiR06, and then permeabilized by digitonin. Oxygen consumption was evaluated using the Oxygraph-2k instrument for Complex I/II-linked respiration in the presence of the proper substrates and inhibitors of the other complex. The results were normalized to the cell numbers.

### Immunoblotting

Immunoblotting was performed as described previously (54), and target proteins were probed with antibodies listed in Table S1. The Super Signal chemiluminescent ECL-plus (Amersham) was used for detecting target proteins.

### Reverse transcriptase–quantitative PCR

Gene expression analysis using reverse-transcriptase quantitative PCR (RT-qPCR) was performed using the LightCycler 480 (Roche, Basel, Switzerland) as described previously (55). The list of primers used is listed in Table S2.

### *In vivo* xenografts and tumor growth analysis

All experiments were conducted in accordance to the guidelines of the QIMR Berghofer Medical Research Institute Animal Ethics Committee. For the human cell line MDA-MB-231 xenograft model, 3 x 10^6^ cells were prepared in 50% growth factor-reduced Matrigel (BD, Biosciences, Bedford, USA)/PBS and injected into the right 4^th^ inguinal mammary fat pad of 6-week old female immunocompromized NOD/SCID mice. For the murine 4T1.2 and 4T1BR4 syngeneic models, 10^5^ cells prepared in PBS were injected into the right 4^th^ inguinal mammary fat pad of 6-week old female immunocompetent Balb/c mice. For the PDX model, 2 mm^3^ tumor biopsy of the TNBC patient (coded TNBC0019) was implanted into the right 4^th^ inguinal mammary fat pad of 6-week old female immunocompromized NOD/SCID mice. Passaging of PDX tumors of breast cancer was conducted according to the IRB guidelines of the Chang Gung Memorial Hospital, Taoyuan, Taiwan. Once tumor size reached ∼30-50 mm^3^, mice were treated with the vehicle or Mzb given by intra-peritoneal (i.p.) injection (0.15 mg/kg, twice per week) for 2 weeks. Tumor growth was measured three times per week using a digital caliper. To calculate the tumor volume the following formula was used: tumor volume = [Lx W^2^]/2, where W = width of the tumor and L = length of the tumor.

For Mzb and 2-DG combination therapy, mice were treated with either vehicle, Mzb (half-MTD dose of 0.075 mg/kg, twice per week), 2-DG (400 mg/kg, IP, 3 times per week), and the combination of the drugs for 2 weeks. For Mzb and doxorubicin combination therapy, mice were treated with the vehicle, Mzb (0.075 mg/kg, twice per week), doxorubicin (10 mg/kg, IP, once per week), and the combination of the drugs for 2 weeks.

### Flow cytometry analysis of ALDH1^+^ stem-cell population

PDX tumors were mashed, filtered through a 100 μm mesh, and washed with PBS. After washing in RPMI medium, live cells in the interphase were collected from Ficoll solution and treated with DNase I (0.02 mg/ml), hyaluoronidase type V (0.2 mg/ml), collagenase 4 (0.5 mg/ml) at 37°C for 2 h. Cells were washed in PBS with 5% FBS and subjected to RBC lysis before single cell suspensions were obtained from the TryPLE solution (Gibco). ALDH1-positive populations were determined by conducting the ALDE FLUOR™ assay following manufacturers’ instructions (StemCell Technologies, MA, USA). Data acquisition and analysis were performed using the NovoCyte Flow Cytometer (ACEA Biosciences, CA, USA).

### Analysis of metastases

*In vivo* metastasis experiments were conducted in accordance to the guidelines of the QIMR Berghofer Animal Ethics Committee. For murine 4T1BR4 syngeneic model, 10^5^ cells were suspended in PBS and injected into the 4^th^ inguinal mammary fat pad of 6-week old female Balb/c mice. Once tumor size reached ∼250-300 mm^3^, the primary tumors were resected. Two days after the tumor resection, mice were treated with the vehicle or Mzb (0.15 mg/kg, twice per week, i.p.) for 2 weeks. At the end of the treatment, lungs and the brain were excised. For lung metastasis, lungs were perfused with PBS to remove the residual blood. Both lungs and brain were fixed with 10% formalin and stained with H&E to facilitate counting of the micro- and macro-metastatic nodules.

### Isolation of circulating tumor cells

Post 1-week drug treatment, mice were euthanized and 100 µl of blood collected from the left ventricle into Eppendorf tubes containing 100 µl of heparin solution and kept on ice. The samples were centrifuged at 400 x g for 5 min at 4°C. The serum was discarded, and the cell pellet re-suspended in 1 ml of the red blood cell lysis buffer (150 mM NH4Cl, 10 mM KHCO3, and 0.1 mM EDTA) before centrifugation at 400 x g for 5 min at 4°C. The supernatant was discarded, and the cell pellet was re-suspended with 1 ml of cold PBS before subjection to several centrifugation cycles to remove the debris and hemoglobin from tumor cells and lymphocytes. The resulting pellet was resuspended in 500 µl of complete growth media and 50 µl of the tumor cell suspension was transferred into 60 mm culture dishes and incubated for 14 days. CTC colonies that formed after 14 days were fixed and stained with crystal violet, followed by imaging and by de-staining for relative quantification.

### Statistical analysis

All values are presented as mean **±** SEM. Data were analyzed using GraphPad Prism 6 (GraphPad Software, CA, USA). Statistical significance was determined by One-way ANOVA followed by Tukey’s post-test and Two-way ANOVA followed by Sidak’s post-test. All the data are expressed as mean values±SEM. Where applicable, statistical significance is denoted by * for P≤0.05, ** for P≤0.01, *** for P≤0.001, and **** for P≤0.0001.

### Data and materials availability

4T1BR4 cells were obtained from Dr Normand Pouliot under the material transfer agreement. The mass spectrometry proteomics data have been deposited to the Proteome Xchange Consortium via the PRIDE partner repository with the dataset identifier PXD015141. The request for any material should be directed to Prof Kum Kum Khanna.

## Acknowledgments

We thank the personnel of the QIMRB Animal, Histology, and Proteomic facilities for their expert technical assistance. We also thank our funding bodies. PR is supported by Cure Cancer Australia & Can Too Foundation Project Grant [ID 1164443], KKK is supported by the National Health & Medical Research Council (NH&MRC) Program Grant [ID 1017028] and the National Breast Cancer Foundation grant [IIRS-19-080], JN was supported in part by the Australian Research Council Project Grant [ID DP180103426] and by the Agency of Health Research Grant [ID 17-30138A].

## Author contributions

PR contributed in study concept and design, acquisition of data, analysis and interpretation of data. PR, KKK, JN, and MK contributed in experimental design, analysis, interpretation of data, and in writing of the manuscript. AL carried out the PDX work. DS contributed in conducting animal experiments. KKD, HG, and MD contributed in proteomic experiments and data analysis. LFD performed respiration experiments. XL contributed in RT-qPCR experiments. PKC analyzed the clinical data. KKK and MK supervised PR in this study. All authors contributed to review and revision of this manuscript.

## Conflict of interests

The authors have declared that no conflict of interest exists.

