## Supplementary file for "Marizomib suppresses triple-negative breast cancer via proteasome and oxidative phosphorylation inhibition"

### Supplementary Files:

### Materials & Methods:

**Proteasome activity assays:**

The CT-L, T-L, and CP-L proteasome activities were measured using the Proteasome-Glo™ Assays Kit (Promega) as per the manufacturer's guidelines.

**Cell viability assays:**

The breast cancer cells and non-malignant mammary epithelial cells were treated with Mzb for 6 days in 48-wells plate, and cell viability was analyzed by the CellTiter 96® AQueous One Solution Cell Proliferation Assay (MTS) (Promega) as per the manufacturer's guidelines.

**Colony formation assays:**

The cells were treated with or without Mzb for 24 h and the effect of Mzb on a long-term colony formation capacity was analyzed as described previously[1].

**3D spheroids assays:**

The 3D tumor spheroid assays were performed using well established techniques as described previously[2]. At the end of the treatment, images of tumor spheroids were taken by EVOS® Digital Inverted Microscope (AMG, WA, USA).

**Caspase-3 activity assays:**

Caspase-3 activity within the treated and untreated breast cancer cells and non-malignant mammary epithelial cells was determined as described previously following the cleavage of Ac-DEVD-AMC (Enzo Life Sciences, NY, USA), a caspase-3 substrate[3].

**ATP, ROS, and intracellular lactate measurement:**

For ATP levels, cells were treated with Mzb for 24 h in 96-wells plate, and intracellular ATP was determined using the CellTiter-Glo® Luminescent assay (Promega). For ROS levels, cells were treated with Mzb for 24 h in 96-wells plate, and intracellular ROS levels were analyzed by the ROS-Glo™ $H_2O_2$ assay kit (Promega). For assessing the intracellular lactate, cells were treated with Mzb for 16 h in 96-wells plate and intracellular lactate levels were analyzed by the Lactate-Glo™ assay (Promega).

**Plasmid DNA transfection:**

SUM159PT cells were transfected with either pcDNA4 empty vector or pcDNA4-Myc-PGC-1α plasmid using Lipofectamine 3000 (Promega) as per the manufacturer's guidelines. The pcDNA4-myc-PGC-1α was a gift from Toren Finkel (Addgene plasmid # 10974; http://n2t.net/addgene:10974; RRID: Addgene_10974)[4].

***In vitro* migration assays:**

The migration assays were performed using Corning® Transwell Inserts. Cells were treated with or without 100 nM Mzb for 24 h and washed. 1 X $10^5$ cells were re-suspended in 0.1% FBS containing DMEM media and were seeded in the upper compartment of the inserts. In the bottom compartment, 10% FBS containing DMEM media was placed as a chemo-attractant. Cells were incubated for 24 h and the migrated cells were fixed with 0.05% crystal violet for 30 minutes and imaged.

**Immunohistochemistry:**

Immunohistochemical analysis was performed using primary 4T1.2 tumors as described previously[5]. ApopTag staining was performed using the ApopTag peroxidase *in situ* apoptosis detection kit (S7100; Millipore-Sigma, Billerica, MA, USA).

**Proteomic analysis:**

All chemicals and solvents are of LCMS grade and were purchased from Merck, USA, unless stated otherwise. All plastic consumables were purchased from Eppendorf, Germany.

*1. In-solution protein digestion*

Total protein was extracted using 2%SDS protein lysis buffer. Sample protein concentrations were determined using the Pierce BCA protein assay kit (Thermo Fischer, USA), following the kit instructions. 30 µg of each protein sample was subjected to reduction, alkylation and trypsin-protein precipitation, prior to in-solution digestion. Briefly, the native disulfide bonds in the protein samples were sequentially reduced and alkylated with 10 mM tris (2-carboxyethyl) phosphine (60°C,30 min), and 40 mM 2-chloroacetamide (dark, RT, 30 min) respectively. Following this, 10 volumes of chilled 100% methanol was added to the sample to facilitate protein co-precipitation along with trypsin (1:100 enzyme: protein; sequencing grade porcine, Promega, USA). After incubating at -20°C for 24 h, samples were centrifuged at 16,000 g for 15 min at 4 °C to collect the protein pellet. The pellet was washed multiple times with chilled methanol to remove residual detergent from the lysis buffer. Trypsin (1:100, enzyme: protein) is added to the protein pellet and then the tube was incubated at 37°C for 18 h. After acidification to a final concentration of 1% v/v formic acid (FA); samples were desalted on the Strata X-33 µm reverse phase solid phase extraction resin (Phenomenex, USA), dried down and resuspended in 12 µl of 0.1 % v/v triflouoroacetic acid (TFA). Prior to LC-MS/MS, the digested peptide concentration was determined using the Pierce µBCA protein assay kit (Thermo Fischer, USA), following the kit instructions.

*2. Tandem liquid chromatography - mass spectrometry (LC-MS/MS) and database search.*

One µg digested peptides were analyzed using a Thermo Scientific VelosPro Orbitrap mass spectrometer (Thermo Fisher, USA) coupled to a Shimadzu Prominence Nano HPLC (Shimadzu, Kyoto, Japan). 5 µl of injected samples were separated on a ProteCol C18 analytical column [(150 mm x 150 µm, 3 µm; connected to a ProteCol guard column, 10 mm x 300 µm, 3 µm), Trajan Bioscience, Australia] over a gradient of 120 min at a flow rate of 1 µl / min. The peptides were eluted using Buffer A (0.1% v/v FA in water) and Buffer B (80% v/v acetonitrile in 0.1% v/v FA), over the specified gradient for 120 min (5.5% B to 40% B at 100 min). The column was maintained at 45 °C. Data acquisition on the mass spectrometer was done using the Data Dependent Top 15 method. The MS spectra were acquired in the mass range = m/z 380 — 1700 (Orbitrap resolution = 60000). Fragmentation for the MS/MS spectra were acquired using collision induced dissociation (CID) in the ion trap mode (dynamic exclusion was set at 90.00 sec).

The extracted raw data was searched for protein IDs against the reviewed human proteome database (49,070 Swissprot entries; database accessed on 01/07/2018) using MaxQuant[1] software, v. 1.5.8.3. MaxQuant parameters were set as follows: digestion = trypsin, with 2 missed cleavages; fixed modification was set to carbamidomethyl; variable modifications = none; LFQ= enabled with minimum ratio count set to 2; match between runs = true; unique and razor peptides were used for protein identification, with minimum unique peptide = 1. The generated protein list was manually filtered to remove contaminants and reverse identified protein IDs; a cut-off score of 2 for minimum peptide counts per protein was applied. Summed intensity-based normalization was carried out. A fold change (FC) cut-off of 1.5 was used to identify differentially regulated proteins (Overexpressed = log2FC≥0.6; Downregulated = log2FC≤0.6). Functional annotation and pathway enrichment analysis of differentially regulated proteins was performed using default parameters of DAVID[6]. The mass spectrometry proteomics data have been deposited to the ProteomeXchange Consortium via the PRIDE[7] partner repository with the dataset identifier PXD015141.

**Supplementary Figures:**

**Fig S1**

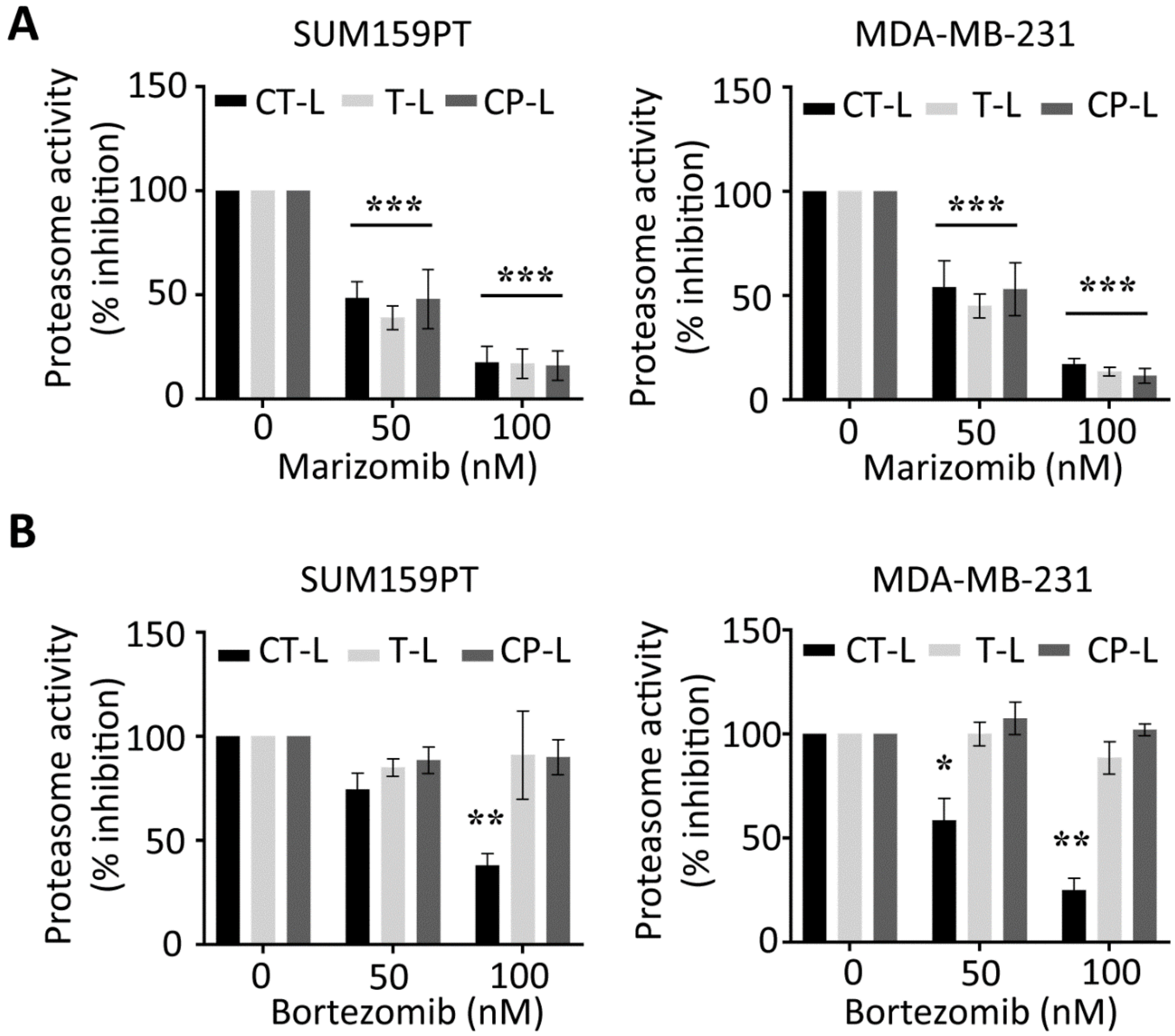

**Fig S1: Marizomib inhibits CT-L, T-L, and CP-L proteasome activity in TNBC cells.**

**(A, B**) SUM159PT and MDA-MB-231 were treated with indicated concentrations of marizomib (0-100 nM) (A) and bortezomib (0-100 nM) (B) for 24 h. The chemotrypsin-like (CT-L), trypsin-like (T-L), and caspase-like (CP-L) proteasome activities were analyzed using Proteasome-Glo® Assays. The activity was calculated relative to untreated control.

#### Fig S2

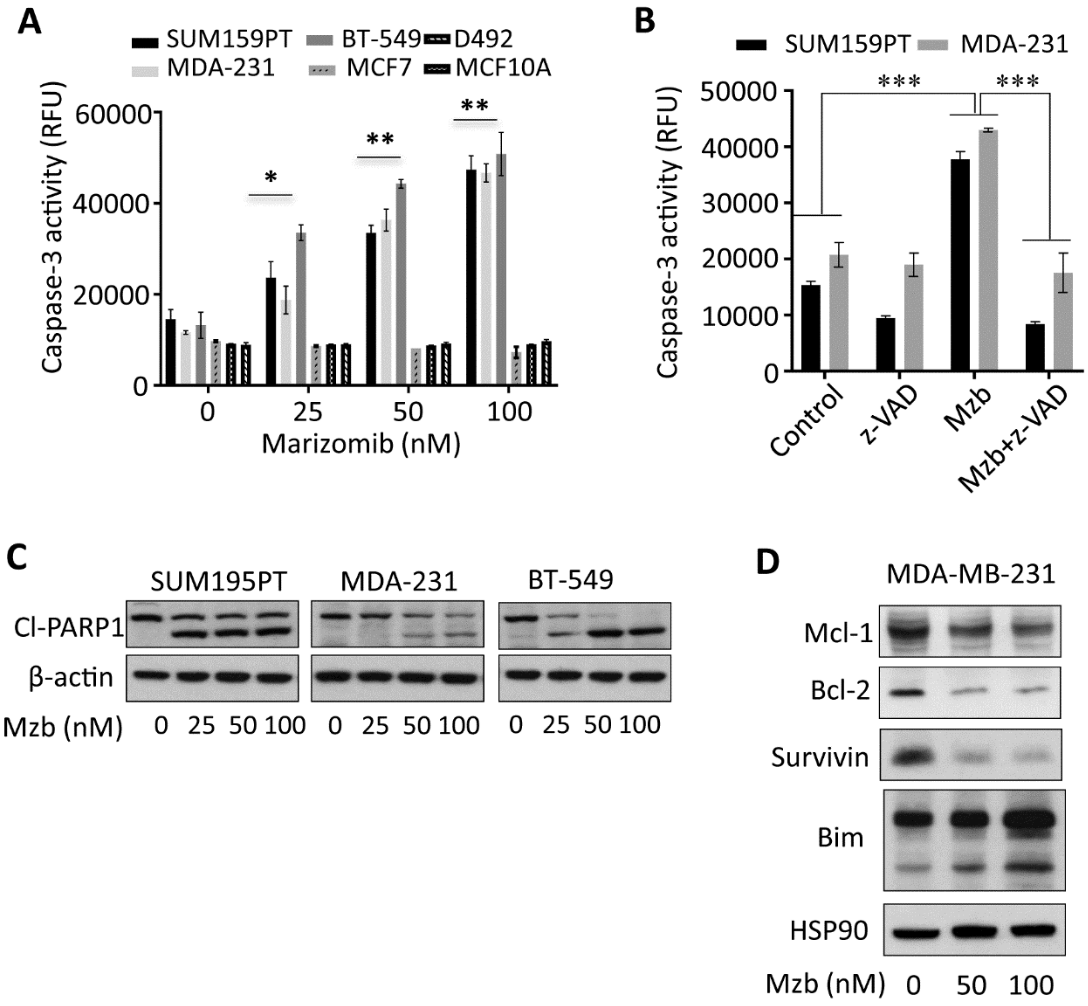

**Fig S2: Mzb induces caspase-3-dependent apoptosis in TNBC cells.**

**(A)** Indicated TNBC, non-TNBCs, and non-malignant cell lines were treated with Mzb (0-100 nM) for 24 h. Caspase-3 activity was analyzed by measuring the cleavage of Caspase-3-specific substrate Ac-DEVD-AMC.

**(B)** SUM159PT and MDA-MB-231 cells were pre-treated with a pan-Caspase inhibitor z-VAD-FMC (50 µM) for 2 h, and subsequently treated with or without Mzb (100 nM) for additional 24

h. Caspase-3 activity was determined by measuring Ac-DEVD-AMC cleavage. One-way ANOVA followed by Tukey's post-tests were employed.

**(C**) TNBC lines (SUM159PT, MDA-MB-231, and BT-549) and near-normal mammary epithelial lines (MCF10A and D492) were treated with Mzb (0-100 nM) for 24 h. The protein levels of cleaved PARP1 were analyzed by western blotting. β-actin was used as a loading control.

**(D)** MDA-MB-231 cells were treated with Mzb (0-100 nM) for 24 h and protein levels of indicated anti- and pro-apoptotic proteins were analyzed by western blotting. HSP90 was used as a loading control.

**Fig S3**

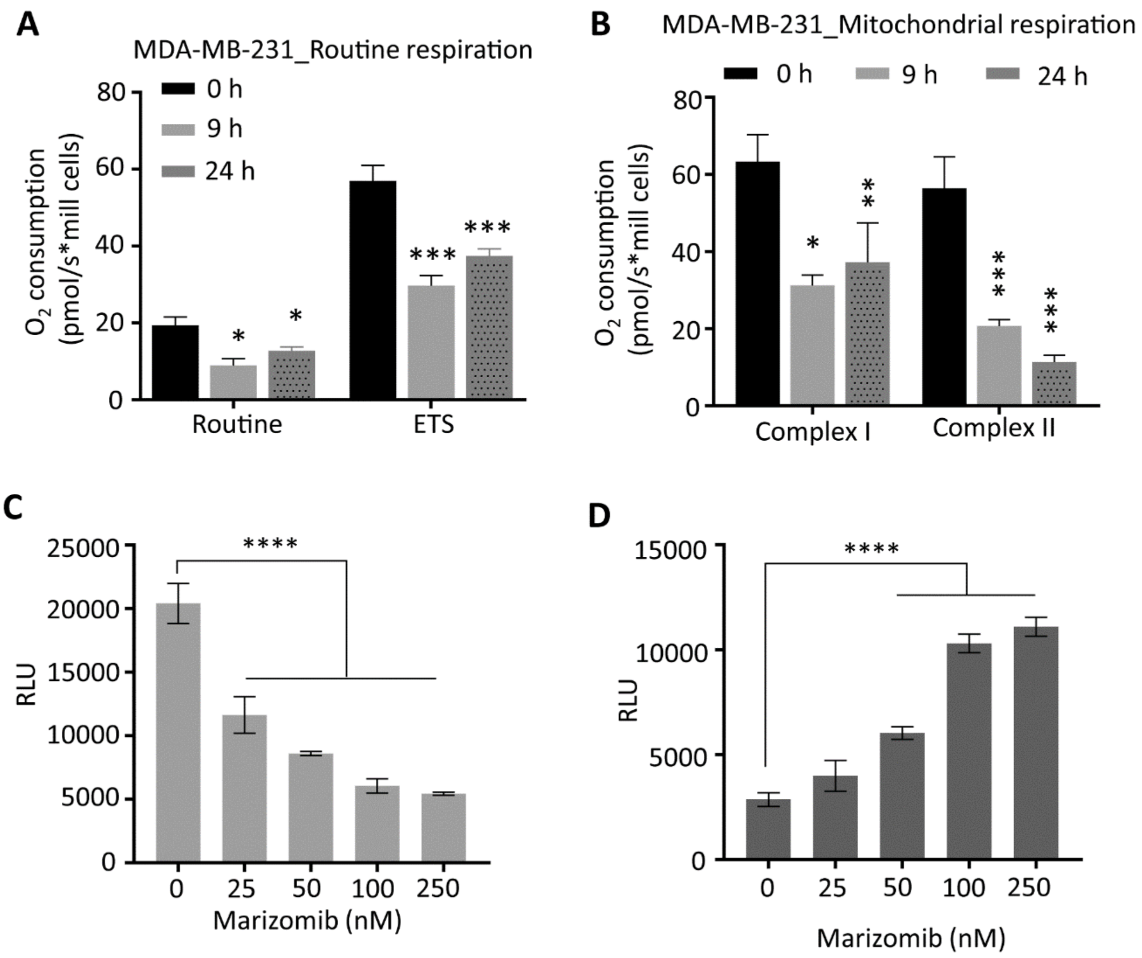

**Fig S3: Marizomib inhibits OXPHOS in TNBC cells.**

**(A)** MDA-MB-231 cells were treated with Mzb (100 nM) for 9 h and 24 h. Oxygen consumption for ROUTINE respiration and FCCP-stimulated uncoupled respiration capacity (ETS) were evaluated on intact cells (n=4 or 5).

**(B)** MDA-MB-231 cells were treated with Mzb (100 nM) for 9 h and 24 h. Oxygen consumption of the cells was evaluated for Complex I/CII-linked respiration (n=4 or 5).

**(C)** MDA-MB-231 cells were treated with Mzb (0-100 nM) for 24 h and intracellular ATP and ROS levels were analyzed. One-way ANOVA followed by Tukey’s post-tests were employed.

**Fig S4**

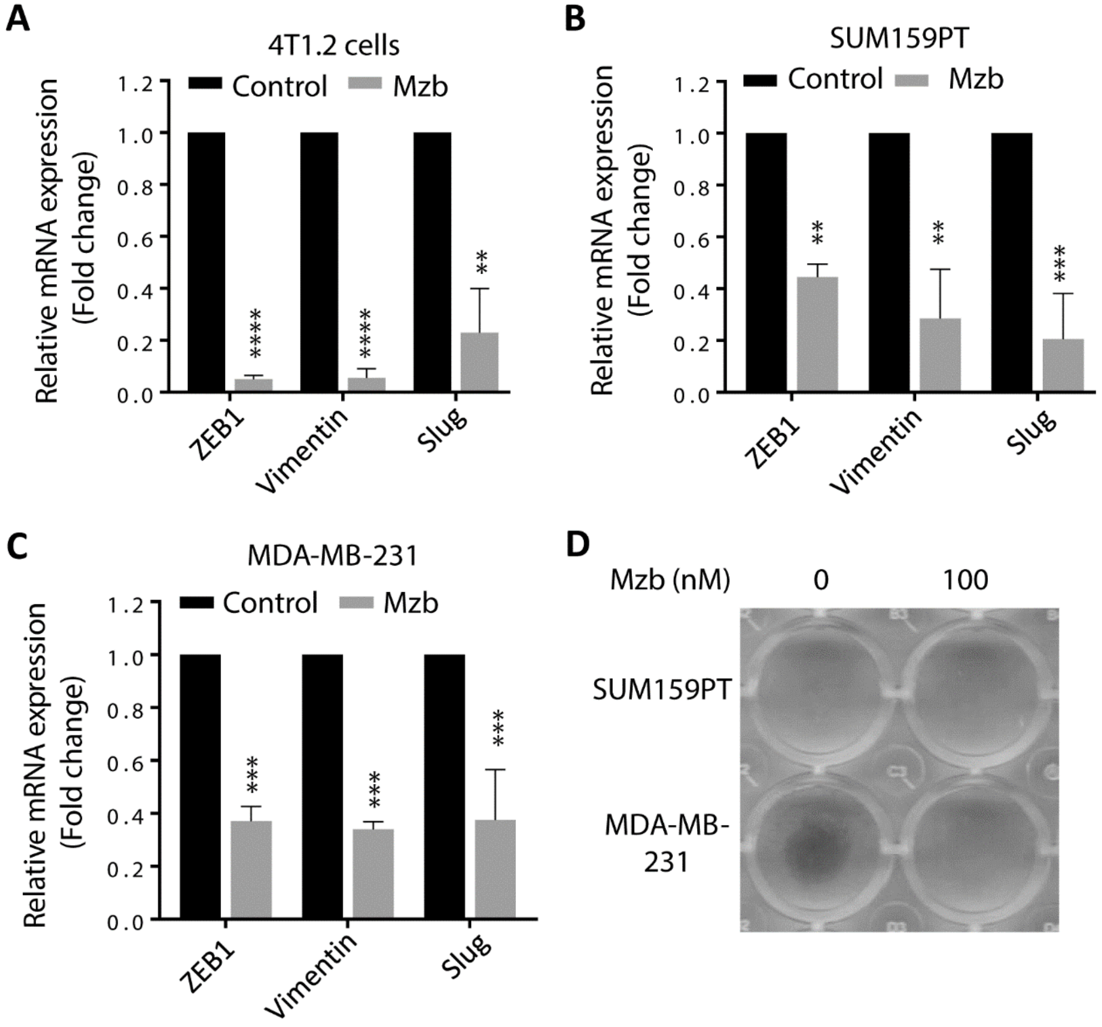

**Fig S4: Marizomib reduces the expression of EMT markers and TNBC cell migration:**

**(A-C)** 4T1.2 (A), SUM159PT (B), and MDA-MB-231 (C) cells were treated with Mzb (100 nM) for 8 h, and mRNA levels of ZEB1, Vimentin, and Slug were analyzed by RT-qPCR. The unpaired "*t*" test was performed.

**(D)** SUM159PT and MDA-MB-231 cells were treated with Mzb (100 nM) for 4 h, and cell migration was analyzed by the Transwell migration assay. Representative images of two independent experiments are shown.

#### Supplementary Table S1:

| Antibody | Supplier | Catalogue number |
|---|---|---|
| Anti-Mcl-1 | Abcam | ab32087 |
| Anti-Bcl-2 | Cell signaling technology | 3498T |
| Anti-Bim | Cell signaling technology | 2933T |
| Survivin | Cell signaling technology | 2808T |
| Anti-β-actin | BD Transduction Laboratories | 612656 |
| Anti-PAPR1 | Cell signaling technology | 9542S |
| Anti-OXPHOS cocktail | Abcam | ab110411 |
| Anti-PGC-1α | Cell signaling technology | 2178S |
| Anti-HSP90 | Santa Cruz | sc-69703 |

**Supplementary Table S2:**

| Gene name | Forward primer | Reverse primer |
|---|---|---|
| Ms PGC-1α | ACGTCCCTGCTCAGAGCTT | CCTTGGGGTCATTTGGTGAC |
| Ms ZEB1 | CAACAAGACACCGCCGTCAT | GAGCAGCTGAAGTTGTCCTC |
| Ms Vimentin | GGATCAGCTCACCAACGACA | CTGCAGCTCCTGGATCTCTT |
| Ms Slug | TGTCTGCAAGATCTGTGGCAA | GAAGCGACATTCTGGAGAAGG |
| Hu PGC-1α | CCTGCATGAGTGTGTGCTCT | TGGGGTCATTTGGTGACTCTG |
| Hu SDHA | GGATGTCGTGGAGAGGGAGGCATT | GGTGCAGCTGCAGGTAGACGTGAT |
| Hu SDHB | TAGCACCAGCTGCCCCAGCTACT | CCCTGGATTCAGACCCTTAGGACA |
| Hu NDUFB8 | CCCCACACCTGTTTCTTGGCATGT | CCACGAAGCCTCCTCAGATCTCAT |
| Hu NDUFB4 | TCGACCCAGCCGAATACAAC | GCAGGATTTTCGATGAGCCC |
| Hu RPL32 | CAGGGTTCGTAGAAGATTCAAGGG | CTTGGAGGAAAACATTGTGAGCGATC |
| Hu ZEB1 | GCAGCTGACTGTGAAGGTGT | CTGTACATCCTGCTTCATCTG |
| Hu Vimentin | CGTGTATGCCACGCGCTCCT | TCGAGCTCGGCCAGCAGGAT |
| Hu Slug | GCACATCCGAAGCCACAC | GGAGAAGGTCCGAGCACAC |
